# Contextualizing Models: Deriving a Kinetic Model of Cancer Metabolism including Growth via Stoichiometric Reduction

**DOI:** 10.1101/2025.08.29.672875

**Authors:** Daria Komkova, Mareike Simon, Jana Wolf, Ralf Steuer

## Abstract

Genome-scale metabolic models (GEMs) offer unprecedented possibilities to study human metabolism, including alterations in cancers. Yet, analyses of GEMs still entail several disadvantages. In particular, constraint-based methods, such as flux balance analysis, are typically restricted to analyse steady-state flux distributions. In contrast, kinetic models based on ordinary differential equations allow assessment of regulatory properties and dynamics. Building kinetic models, however, is still hampered by the lack of knowledge about kinetic parameters and is typically focused on individual pathways.

Here, we present an approach to derive kinetic models of metabolism augmented by coarse-grained overall reactions that represent the remaining cellular metabolism and biosynthetic processes. Using algorithmic network reduction, we derive coarse-grained reactions that preserve the correct stoichiometry of precursors, energy, and redox equivalents required for cellular growth. Analysis of the GEM-embedded kinetic model uses Monte Carlo sampling to address parameter uncertainty. We exemplify our approach by constructing a kinetic model of cancer metabolism that includes an explicit description of cellular growth. We show that the GEM-embedded kinetic model differs in its control properties from the corresponding model without growth, with implications for understanding regulatory hotspots and drug target identification.

## Introduction

Metabolic reprogramming to support enhanced cell growth and proliferation is widely recognized as a hallmark of cancer (Hanahan and Weinberg, 2011), and understanding cancer-related metabolic alterations has attracted substantial efforts over the past decade (DeBerardinis and Chandel, 2016; Pavlova and Thompson, 2016; Vander Heiden and DeBerardinis, 2017). Concomitant to advances in analytical tools, computational models of cancer metabolism, have played a crucial role to investigate the consequences of metabolic reprogramming and to identify potential therapeutic interventions and possible drug targets (Ghaffari et al, 2015b; Yizhak et al, 2015; Resendis-Antonio et al, 2015). In particular, genome-scale metabolic models (GEMs) provide a computational framework to analyze metabolic phenotypes, facilitate multi-omics data integration, and investigate metabolic reprogramming in cancer cells (Folger et al, 2011; Shlomi et al, 2011; Agren et al, 2014; Ghaffari et al, 2015b). GEMs seek to provide a comprehensive account of the stoichiometry of all possible metabolic interconversions in a tissue or cell by establishing gene-enzyme-reaction relationships derived from the annotated genome sequence, literature data, and reaction databases (Kanehisa and Goto, 2000; Milacic et al, 2023). Following the publication of the human genome (Lander et al, 2001), the first human GEMs were presented in 2007 (Duarte et al, 2007), with continuous updates and improvements resulting in the latest iteration, the human metabolic atlas (Robinson et al, 2020). A human genome-derived GEM allows researchers to extract context-, tissue- or cell-specific subnetworks, based on transcriptomic or proteomic data. The analysis of context-specific GEMs is widely applied to the analysis of cancer metabolism (Jerby et al, 2010; Agren et al, 2012; Ghaffari et al, 2015a; Lewis et al, 2021).

Notwithstanding its merits, however, the analysis of GEMs using constraint-based methods also entails several drawbacks and disadvantages. In particular, constraint-based methods, such as flux balance analysis (FBA), are typically restricted to steady-state properties of metabolic fluxes derived from the stoichiometric matrix. In addition, since solutions obtained from constraints on the mass balance of the stoichiometric matrix are typically under-determined, applications require the definition of a suitable objective function, such as the maximization of biomass yield given constraints on uptake fluxes, to predict relevant metabolic properties.

In contrast, kinetic models of metabolism, based on ordinary differential equations (ODEs), allow us to study the dynamics and regulatory properties (Teusink et al, 2000; Wolf et al, 2000; Wolf and Heinrich, 2000; Steuer and Junker, 2009; Murabito et al, 2011; Roy and Finley, 2017; Shestov et al, 2014; Mulukutla et al, 2015; Marín-Hernández et al, 2011; Marín-Hernández et al, 2014; Moreno-Sánchez et al, 2018; Snaebjornsson et al, 2025). The construction of kinetic models, however, requires detailed knowledge about the biochemical processes, such as appropriate kinetic rate equations and enzyme-kinetic parameters for each reaction–knowledge that is typically available only for a small set of reactions. Hence, the construction of kinetic models is currently not straightforwardly feasible on a genome-scale and is typically focused on individual pathways or specific parts of metabolism. As a consequence, kinetic models often do not incorporate the metabolic context of the respective pathway, such as withdrawal of metabolic intermediates and energy and redox equivalents required for growth and other cellular functions. Instead, adjacent energy-consuming pathways are often modelled using a general ATP hydrolysis reaction (ATPase), sometimes, in particular in models of glycolysis, as the only intracellular metabolic sink. In models where a description of cellular growth is required, the growth rate has previously been implemented in dependence of internal metabolites concentrations, such as ATP and glutamine, without necessarily accounting for the withdrawal of metabolic intermediates, energy and redox equivalents in the correct stoichiometric ratios.

To overcome these limitations, and to put pathway-focussed kinetic models into a broader whole-cell context, we seek to implement a workflow that augments a kinetic model of core cancer metabolism with stoichiometrically correct coarse-grained reactions to describe cellular growth. Specifically, we make use of stoichiometric network reduction (Erdrich et al, 2015; Röhl and Bockmayr, 2017; Singh and Lercher, 2020) to derive a reduced stoichiometric network that describes the metabolism of a cancer cell line. The reduction algorithms allows us to retain a set of core reactions, including glucose uptake and the glycolytic pathway, whereas the remaining reactions are aggregated into coarse-grained overall reactions. These coarse-grained overall reactions are stoichiometrically consistent with the full model and its requirements for growth. That is, the coarse-grained reactions withdraw metabolic intermediates, energy and redox equivalents in the correct stoichiometric ratios and amounts required for growth.

We discuss and exemplify our workflow using a model of aerobic cancer glycolysis previously developed by Shestov et al (2014). Our aim is to augment this model with an explicit description of cellular growth, and to study the differences in dynamic properties and the metabolic control profiles. We show that the GEM-embedded model provides an estimate of a biochemically plausible growth rate as a function of nutrient uptake, retains relevant conservation relationships, and exhibits a modified control profile compared to the original model. To account for unknown enzyme-kinetic parameters, we further utilize a Monte-Carlo (MC) approach based on parameter sampling to assess the dependency and robustness of control properties with respect to the choice of enzyme-kinetic parameters. Our approach provides a general template to construct and contextualize kinetic models of metabolism.

## Results and Discussion

### Summary of the workflow

Our approach is visualized in Figure 1. Individual steps are detailed in the following sections. In brief, our starting point is a genome-scale reconstruction of human metabolism, Human1 (Robinson et al, 2020), from which a cell-type and context-specific stoichiometric model is extracted (Agren et al, 2014). The cell-type specific model is then reduced using algorithms for stoichiometric network reduction (Erdrich et al, 2015). Stoichiometric network reduction requires us to specify a minimal set of metabolic functions that the reduced model must be capable of. Here, the specifications are such that the reduced network (i) must allow for an identical rate of biomass formation on a defined medium as the original model, (ii) must be capable of growth-independent ATP production from glucose under aerobic as well as anaerobic conditions, and (iii) must include all reactions of core glycolysis.

**Fig. 1.**
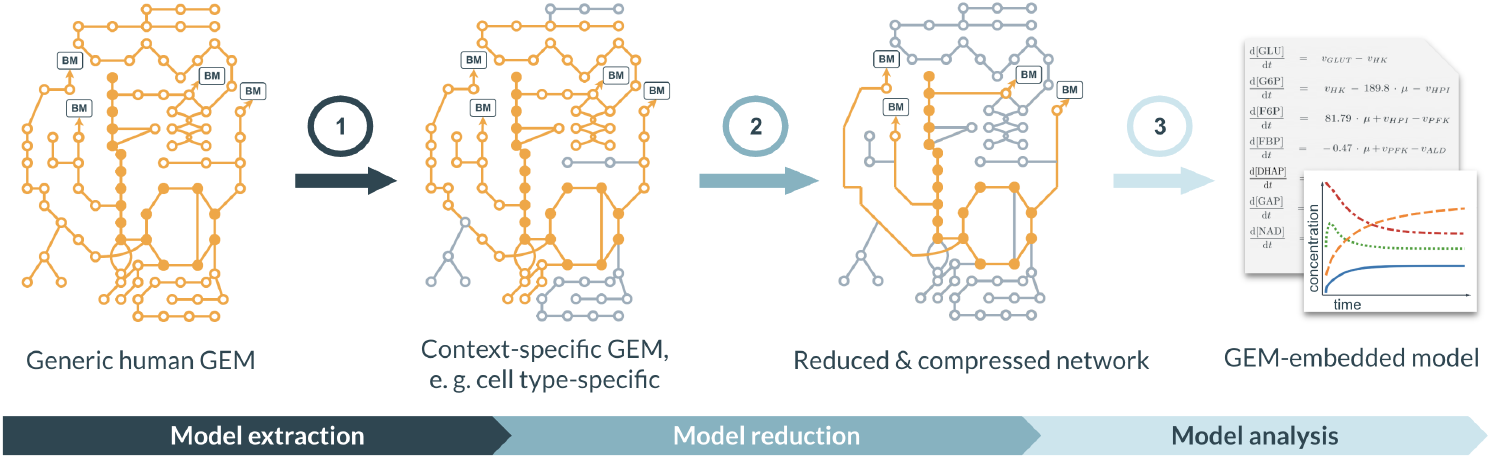
Summary of the workflow. Our aim is to derive a GEM-embedded kinetic model that is capable to describe the kinetics of a metabolic pathway within a cellular context. (1) Starting point is a genome-scale metabolic reconstruction from which a tissue-specific stoichiometric model is extracted. (2) The tissue- or cell type-specific model is subject to stoichiometric network reduction such that desired reactions are retained and the remaining metabolism is aggregated into coarse-grained overall reactions. (3) The reduced stoichiometry is converted into a GEM-embedded kinetic model by assigning kinetic rate equations and parameters to all reactions. The dynamics and control properties of the GEM-embedded kinetic model are then evaluated. The robustness with respect to parameterization is evaluated using parameter sampling.

The reduced model is then subject to lossless compression and further reduced to a GEM-embedded glycolytic core model that consists of 18 reactions, 19 internal metabolites and 4 mass conservation relationships, resulting in 15 independent internal metabolites. The GEM-embedded glycolytic core model includes an explicit description of cellular growth and oxidative phosphorylation, and is evaluated with respect to its stoichiometric and kinetic properties.

Of particular interest are shifts in the control profile of glycolytic flux compared to a previously published model of cancer glycolysis that lacks the metabolic context provided by the genome-scale stoichiometric reduction. To evaluate the robustness of the results, we utilize conditional Monte-Carlo (MC) sampling of kinetic parameters, following established methods (Steuer and Junker, 2009; Murabito et al, 2014). Conditional MC sampling of parameters allows us to evaluate the control properties of the system without requiring detailed knowledge of kinetic parameters.

Our approach establishes a template to derive a GEM-embedded kinetic model from a genome-scale metabolic reconstruction and to analyze its stoichiometric and kinetic properties.

### Establishing a cell type-specific reconstruction

Our first step is to establish a suitable cell type-specific stoichiometric model. For proof of concept, we choose the HT-29 cell line, a fast-growing human colon cancer cell line with a division time of approx. 24 hours in Dulbecco’s Modified Eagle Medium (DMEM) (Petitprez et al, 2013). The cell type-specific reconstruction is obtained from the human metabolic reconstruction Human1 (Robinson et al, 2020), using the tINIT algorithm (Agren et al, 2014). The tINIT algorithm uses RNA-seq data to determine which reactions of the full model are present in the context-specific model, and allows us to specify which metabolic tasks the cell type-specific model must be able to fulfill. For the latter, we choose the 57 tasks proposed by Agren et al (2014) that are essential to cell viability (see Materials and Methods, Section ‘Workflow’). The cell type-specific network consists of 8483 reactions and 3008 unique metabolites in 9 cellular compartments and describes growth of the HT-29 cell line.

We note that the choice of the *in silico* medium is crucial for further analysis. We here use an *in silico* medium consisting of 39 extracellular compounds, and constrain the maximum uptake rates for all compounds provided in the medium according to the limits given by Folger et al (2011). In addition, the medium includes also retinoate, alpha-tocopherol, beta-tocopherol, linoleate, and linolenate, to account for the increased nutrient requirements of Human1 compared to earlier reconstructions. The uptake rate of glucose is constrained to 5.0 mmol*/*gDW*/*h, that of glutamine to 0.5 mmol*/*gDW*/*h. The complete list of compounds and uptake constraints are provided in Supplementary Table S1. The network was tested for feasibility of growth in the defined medium.

To obtain a reference solution for model reduction, we maximize the biomass objective function as a proxy for the growth rate using constraints on all exchange metabolites, given in Supplementary Table S1, as a function of the upper bound of the oxygen (O_2_) uptake rate. Figure 2A shows the maximal growth rate, as well as the lactate and glucose exchange fluxes obtained using parsimonious FBA (pFBA, Lewis et al (2010)). For very low upper bounds on the O_2_ uptake rate, growth is limited because O_2_ is required as a substrate for biosynthesis, independent of its use in respiration. At a critical value (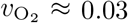 mmol*/*gDW*/*h), the stoichiometric demands for O_2_ for biosynthesis are no longer limiting, and the maximal growth rate is reached. At this point, the ratio of lactate secretion versus glucose uptake (*v*_LACT_*/v*_GLCT_) is at its maximal value (*v*_LACT_*/v*_GLCT_ 1.87).

**Fig. 2.**
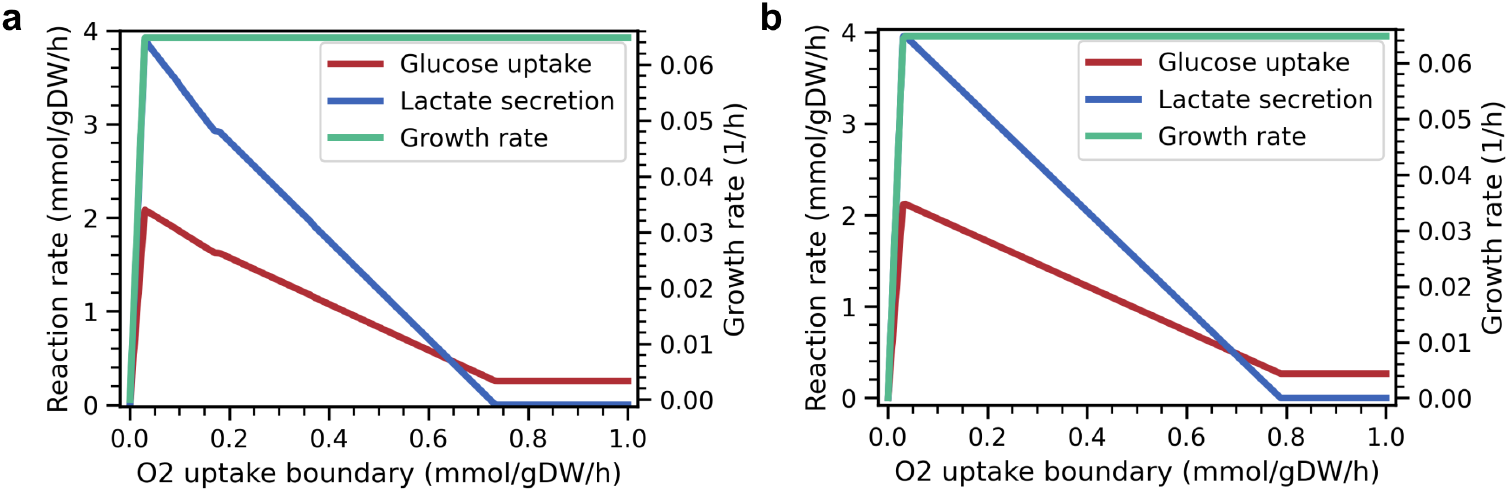
Reference solutions for the cell type-specific model compared to the stoichiometrically reduced model. (A) Growth rate and exchange fluxes (glucose uptake and lactate secretion) for the cell-type specific full stoichiometric model. Lines correspond to a pFBA solution with maximal biomass formation (growth rate) as a function of the constraint on maximal oxygen uptake flux. (B) The corresponding flux solutions after stoichiometric model reduction. The results are not unique, shown is the solution with minimal glucose uptake. For both models, there exists a minimal amount of O_2_ required for growth. Beyond the minimal amount, O_2_ uptake limits respiration and the cell gradually shift from a partially fermentative state to a fully respiratory state. For simplicity, all exchange fluxes are depicted with a positive sign.

For higher upper bounds on the uptake rate, O_2_ is increasingly used as the electron acceptor in respiration and lactate secretion decreases, and hence the ratio *v*_LACT_*/v*_GLCT_, decreases until the cell exclusively relies on oxidative phosphorylation to regenerate ATP (*v*_LACT_*/v*_GLCT_ = 0). We note that, under the constraints chosen here, glucose is not essential for viability and growth, and the maximal growth rate is independent of the upper bound on *v*_O_2. Instead, the metabolites linoleate, linolenate, choline, serine, and aspartate are taken up with maximally allowed rate and therefore are limiting growth. We emphasize that, in the following, we do not analyze a particular FBA solution, the results shown in Figure 2A solely serves to specify a reference solution for stoichiometric model reduction.

### Stoichiometric model reduction

Stoichiometric network reduction is performed using NetworkReducer (Erdrich et al, 2015). The algorithm implements a two-step reduction process. Within the first step, reactions that are not required to support the protected metabolic functions are iteratively removed (pruning). As a second step, stoichiometrically coupled reactions are combined into coarse-grained over-all reactions. The second step does not reduce the feasible flux space of the network (loss free compression), but significantly reduces the number of reactions.

The pruning step requires a list of protected network elements (metabolites and reactions), as well as a list of ‘metabolic functions’ or ‘phenotypes’. The latter are given as FBA or pFBA constraints that the reduced network is required to fulfill. In our case, the protected functions are such that the reduced stoichiometry (i) must allow to regenerate ATP from glucose under aerobic conditions with a yield that matches the full model, (ii) must allow to regenerate ATP from glucose under anaerobic conditions with a yield that matches the full model, and (iii) is capable of growth on the defined medium with a maximal growth rate that matches the growth rate of the full model at the reference point defined by a ratio *v*_LACT_*/v*_GLCT_ = 1.87 of lactate secretion versus glucose consumption (see Fig 2). The protected functions are implemented as a feasibility problem that the reduced network must be capable to fulfill.

In addition to the functional constraints, we protect all reactions of core glycolysis. The resulting reduced stoichiometry, after pruning, consists of 797 reactions and 835 metabolites (including external metabolites) and gives rise to 282 elementary flux modes (EFMs, Schuster and Hilgetag (1994)). The subsequent loss-free compression results in a model with 59 reactions, 46 external metabolites, and 72 internal metabolites subject to 20 mass-conservation relationships, resulting in 52 independent metabolites. Figure 2B shows an FBA solution of the reduced network, which can be directly compared to the solution of the full network in Figure 2A. As shown, the reduction retains the required properties of the full network. We note that the flux solutions are not unique and exhibits considerable variability. Both figures show the solution with minimal glucose uptake. The reduced network is provided as Supplementary File 1 (pruned network, SBML) and Supplementary File 2 (pruned and compressed network, SBML).

While kinetic modeling is, in principle, feasible using the pruned and compressed network, for the purpose of this study we seek to further reduce the network. In particular, we are interested in a direct comparison of the reduced kinetic model with the computational model of cancer glycolysis proposed by Shestov et al (2014). To obtain a stoichiometry similar to Shestov et al (2014), we therefore follow the approaches of Baroukh et al (2014) and Tummler et al (2015) and collapse the EFMs for respiration and biomass formation of the pruned model into two single coarse-grained reactions, using the FBA solution as a reference. The final network is shown in Figure 3 and consists of 18 metabolic reactions and 19 internal metabolites, which are subject to 4 mass conservation relationships. A list of reaction stoichiometries is provided in Table S2 in the Supplementary Information.

**Fig. 3.**
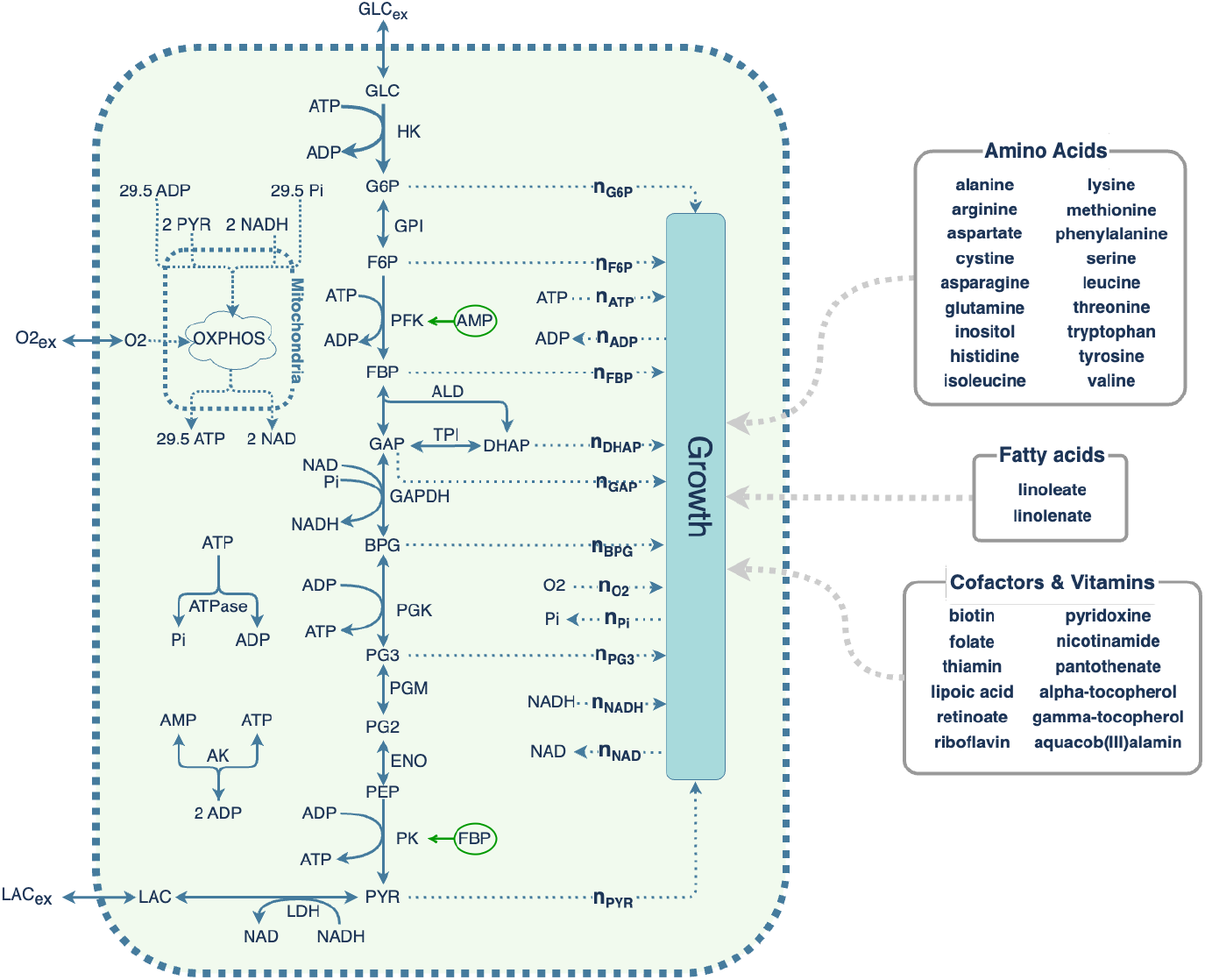
Schematic of the final GEM-embedded model. The model describes glycolysis, the fermentation of glucose to lactate, oxidative phosphorylation, and growth. It consists of 18 reactions, 19 internal metabolites and includes four mass conservation relationships. A list of abbreviations is provided in Materials and Methods. The stoichiometric coefficients of the growth reaction are given in the main text and in Supplementary Table S4. Other nutrients are taken up proportionally to the growth reaction but are not explicitly included in the kinetic model.

### Stoichiometric properties of the GEM-embedded model

Consistent with the objectives of the reduction, the reduced model describes the glycolytic pathway, as well as ATP regeneration and the formation of biomass. The remaining metabolic processes, including the uptake of other nutrients, are aggregated into the coarse-grained overall reaction for growth. The growth reaction describes the conversion of central metabolites into cellular biomass, as shown in Figure 3, with the overall stoichiometry, 0.0007 mmol BPG + 0.4801 mmol PG3 + 0.1953 mmol DHAP + 0.0043 mmol FBP + 0.4712 mmol O2 + 0.1647 mmol F6P + 0.07 mmol GAP + 0.4129 mmol G6P + 2.8958 mmol NADH + 2.415 mmol PYR + 62.9543 mmol ATP → 1 gDW Biomass + 62.9543 mmol ADP + 2.8958 mmol NAD + 64.287 mmol Pi.

Due to the algorithmic reduction, the full mass-balance of growth is preserved in the reduced model. All remaining nutrients, including nutrients not explicitly modelled, are taken up proportional to the growth reaction and can, in principle, be included in a simulation of batch growth. For the present reduction, approximately 20.3% of carbon in the biomass derives from glucose, close to literature values (Hosios et al, 2016).

We highlight two advantages of this algorithmic reduction. Firstly, the stoichiometric coefficients of the coarse-grained growth reaction preserve mass-balance and account for the correct requirements of co-factors, such as ATP and NADH. While kinetic models sometimes include withdrawal reactions from glycolysis (Teusink et al, 2000), or describe growth as a function of metabolic intermediates (Roy and Finley, 2017), such *ad hoc* assignments mostly do not reflect the actual nutrient and energy requirements of growth, and do typically not allow us to derive a quantitative value for the growth rate based on nutrient uptake.

Secondly, the algorithmic reduction preserves the conservation relationships of the pathway. The GEM-embedded model shown in Figure 3 gives rise to 4 conservation relationships, summarized in Table 1. While conservation relationships involving only a few conserved moieties, such as the conservation of the adenylate moiety, can be readily identified and preserved using *ad hoc* withdrawal reactions, the algorithmic reconstruction also preserves more complex conservation relationships. This indicates that the inclusion of withdrawal reactions without genome-scale stoichiometric context are not always suitable for model construction and may result in systematic errors.

**Table 1.** The GEM-embedded model shown in Figure 3 gives rise to four conservation relationships. The values of total concentrations are part of the parameterization and independent of the stoichiometric reduction.

| Moiety | Metabolites | Value [mmol/l] |
| --- | --- | --- |
| phosphate moiety | $[G6P] + [F6P] + 2 [FBP] + [GAP] + [DHAP] + 2 [BPG] + [PG2] + [PG3] + [PEP] + 2 [ATP] + [P] + [ADP]$ | 13.24 |
| adenylate moiety | $[ATP] + [ADP] + [AMP]$ | 3.02 |
| oxidation state | $[PG3] + [PG2] + [BPG] + [PEP] + [PYR] + [NAD]$ | 2.01 |
| redox potential | $[NADH] + [NAD]$ | 0.55 |

### A kinetic model of cancer metabolism

Given the GEM-embedded stoichiometry shown in Figure 3, the next step is to convert the stoichiometric network into a kinetic model by assigning rate equations and kinetic parameters to all reactions. To this end, protected enzymatic reactions, i.e. reactions that are retained from the original stoichiometry and not compressed into coarse-grained overall reactions, can be parameterized following established workflows (Smith et al, 2018; Foster et al, 2019) making use of kinetic parameters obtained from reaction databases and primary literature (Chang et al, 2021; Wittig et al, 2017).

In the following, however, our aim is a direct comparison with the model derived from Shestov et al (2014) (see Materials and Methods for model definitions), hence we assign all rate equations and parameters of reactions present in both models to the rate equations and parameters used in the original model. In brief, Shestov et al (2014) use reversible Michaelis-Menten equations for all reactions, with thermodynamic information sourced from Goldberg et al (2004) and König et al (2012), and Michaelis-Menten parameters assigned to values that correspond to measured intracellular concentrations of the respective metabolites. Coarse-grained reactions are assigned to irreversible Michaelis-Menten kinetics, with *K*_*M*_ parameters assigned to measured intracellular concentrations of the respective metabolites. We later evaluate different choices of parameter values to assess the robustness of the parameterization.

The maximal reaction velocities of coarse-grained reactions are free parameters that do not correspond to the properties of individual enzymes. Hence, the values are assigned according to plausible biological expectations. Specifically, the maximal reaction velocity *v*_max,BM_ of the coarse-grained biomass reaction is assigned such that it does not limit the rate of biomass formation. Instead, the coarse-grained biomass reaction is limited by the availability of metabolic precursors. This choice reflects the assumptions used for the underlying parsimonious FBA problem: the enzymes that constitute the overall growth reaction are expressed such that the growth rate is close to its maximal value and the growth rate is not limited by the expression of anabolic proteins. Figure 4A shows the dependence of the growth rate as a function of *v*_max,BM_ for different values of external glucose. The parameter *v*_max,BM_ is chosen such that the growth rate reaches a plateau and the growth rate is approximately *µ* = 0.347*d*^−1^ for an external concentration of glucose of [*GLC*_*e*_] = 12 mmol/l, corresponding to experimental values reported by Wahl et al (2002). The precise values of *v*_max,BM_ has no major impact on our further analysis.

**Fig. 4.**
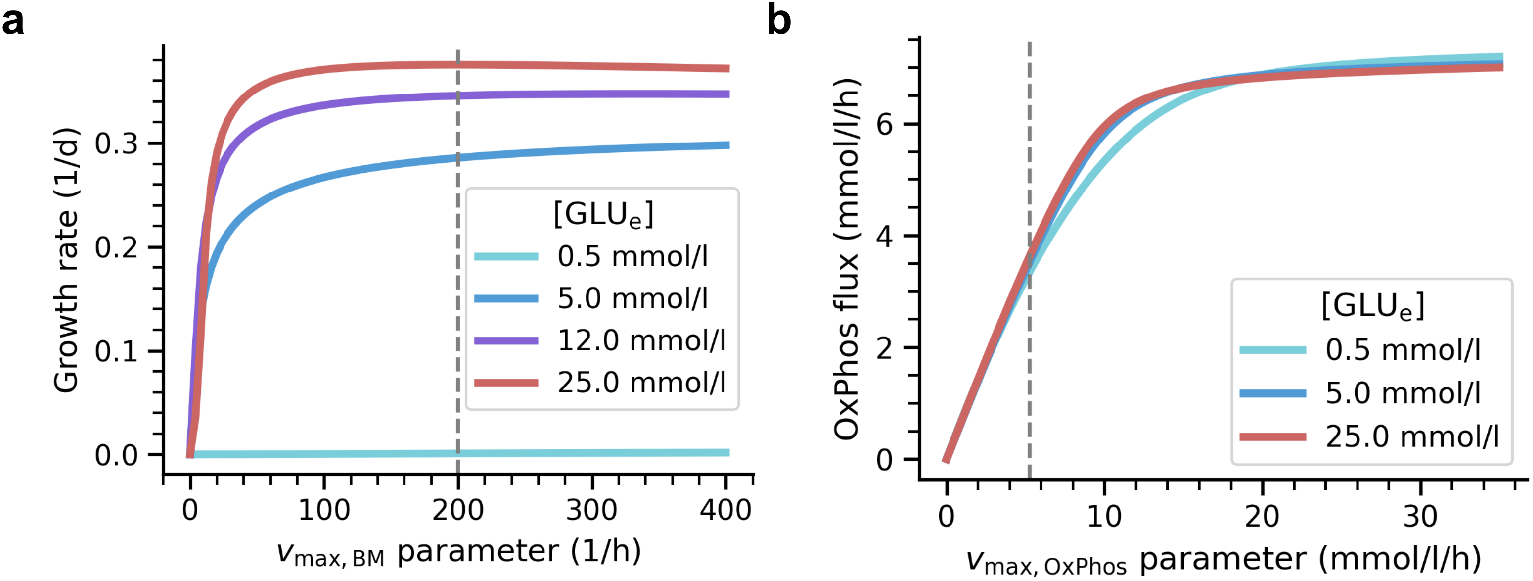
(A) Dependency of the specific growth rate on the maximal reaction velocity *v*_max,BM_ of the coarse-grained overall growth reaction. The parameter is chosen such that growth is mostly saturated (*v*_max,BM_ = 200 1/h, indicated by dashed line). Shown are the results for four concentrations of external glucose. (B) Dependence of OxPhos flux as a function of the maximal reaction velocity *v*_max,OxPhos_ of the coarse-grained OxPhos reaction. The parameter is chosen such that mitochondrial capacity is limiting *v*_max,OxPhos_ = 5.25 mmol/l/h, indicated by dashed line).

Different to the rate of biomass formation, we assume that the mitochondrial capacity, as given by the maximal reaction velocity *v*_max,OxPhos_, is limiting respiration. Hence, we choose *v*_max,OxPhos_ such that the rate of the coarse-grained OxPhos reaction is limited by *v*_max,OxPhos_, as shown in Figure 4B. A list of all kinetic parameters is provided in the Supplemental Materials, Section 3.

### Kinetic properties of the GEM-embedded model

With the GEM-embedded kinetic model fully defined, we now evaluate its kinetic properties in comparison with the model without growth. Firstly, the model allows us to assess the dependency of the predicted growth rate on the external glucose concentration, as well as the dependence of the growth rate on the expression of the glucose transporter (GLCT). Both dependencies are shown in Figure 5. In the following, we choose an external glucose concentration of [*GLC*_*e*_] = 5 mmol/l and a transporter activity *v*_max,GLCT_ = 100 mmol/l/h as a reference point for further analysis.

**Fig. 5.**
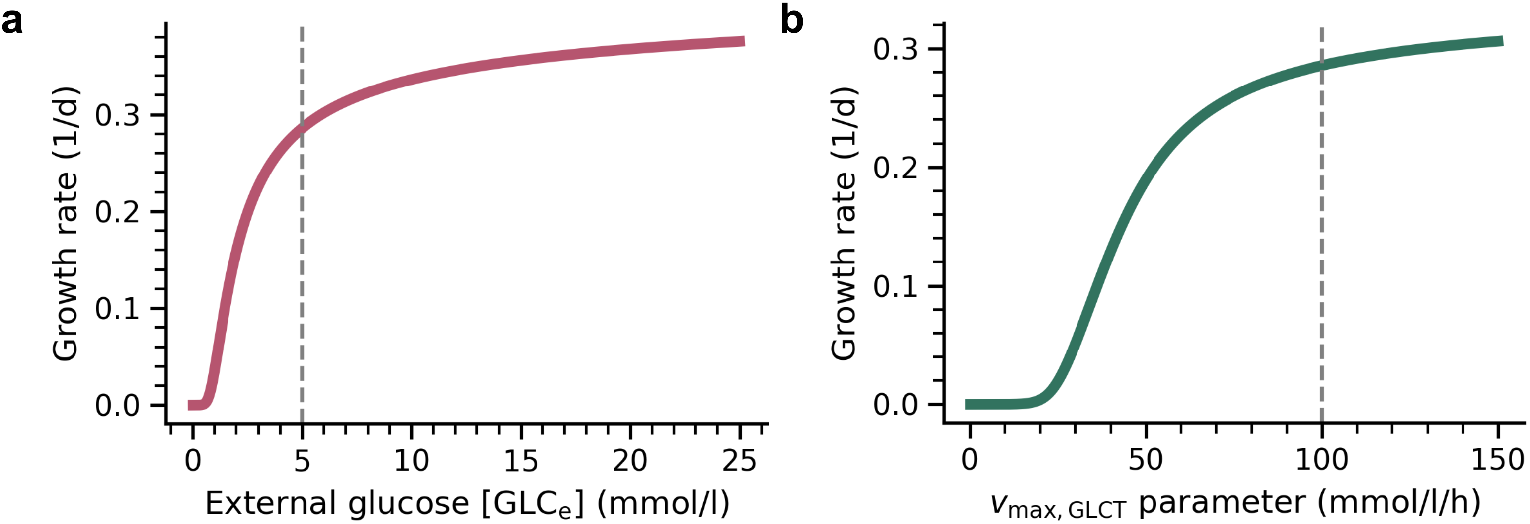
(A) The dependence of the specific growth rate of the GEM-embedded model as a function of the extracellular glucose concentration. The reference value ([*GLCe*] = 5 mmol/l) used in the following is represented by a dashed line. (B) The dependency of the specific growth rate on the activity *v*_max,GLCT_ of the glucose transporter. The dashed line indicates the reference value used in the analysis (*v*_max,GLCT_ = 100 mmol/l/h).

At the reference point, the specific growth rate is *µ*_max_ *≈* 0.29 *d*^−1^, corresponding to a division time of *T*_*D*_ *≈* 57.8 *h*, well within the range of experimental values for cancer cell lines, including the cell line used for the parameterization of the model of Shestov et al. (Wahl et al, 2002). We note that an explicit description of the specific growth rate in dependence of external glucose or expression of transporters is a key feature of GEM-embedded models and not straightforwardly possible in conventional pathway-focused kinetic models.

Next, we assess how the inclusion of the coarse-grained growth reaction affects the distribution of metabolic flux control in the model. To this end, we evaluate the normalized flux control coefficients (FCCs) as defined in metabolic control analysis (MCA, Kacser and Burns (1973); Burns et al (1985); Reder (1988); Steuer and Junker (2009)). The FCCs quantify the response of a metabolic flux to a small change in enzyme expression. A definition is provided in the Materials and Methods.

Figure 6 shows the FCCs of glycolytic enzymes on the hexokinase (HK) reaction, used here as a proxy for glycolytic flux, in comparison to the FCCs obtained from the Shestov-derived model. A positive FCC implies that a small increase in the respective enzyme results in an increase of the HK flux proportional to the value of the FCC. In both models, the control of flux through the HK reaction is distributed across the pathway. We observe differences in the control of both models exerted by upper glycolysis, in particular glucose transport (GLCT), hexokinase (HK)and phosphofructokinase (PFK). Control exerted by enzymes of lower glycolysis is mostly comparable, e.g. high control is exerted by pyruvate kinase (PK) and the lactate transport (LACT). In both models, GAPDH exhibits a high positive control over HK flux.

**Fig. 6.**
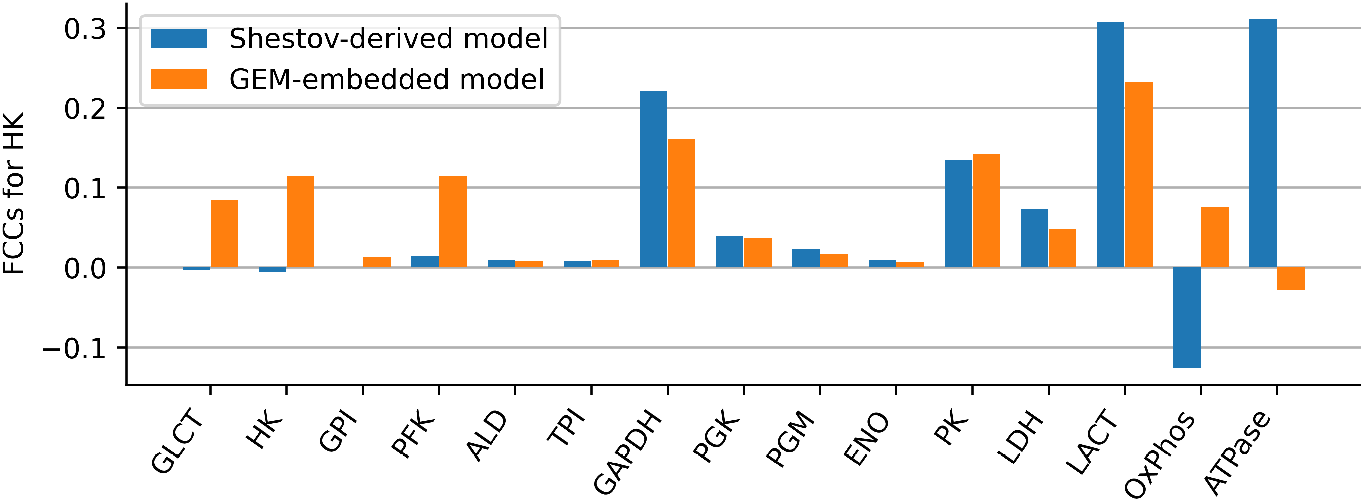
Flux control coefficients (FCCs) on the hexokinase reaction as a proxy for glycolytic flux. Shown are the FCCs for the GEM-embedded model in comparison to the Shestov-derived model. The overall control profile is similar, key differences are discussed in the text. We note that coarse-grained reactions do not correspond to a single enzyme whose expression can be altered. Therefore, for these reactions, the FCCs should be interpreted as an indicator of the sensitivity with respect to the assigned parameter value.

Notably, the FCCs of several reactions, in particular OxPhos and ATPase change signs in the GEM-embedded model. We note that in the model derived from Shestov et al (2014), ATPase is the sole energy sink within the model, implicitly also accounting for ATP used for cellular growth. In contrast, within the GEM-embedded model, the ATPase represents an separate ATP sink (non-growth-associated ATP maintenance, NGAM), in addition to the coarse-grained growth reaction.

Beyond the FCCs for HK, Figure 7 provides a global comparison of FCCs between both models. Shown is the matrix of FCCs for both models. In addition, Figure 8 provides a direct comparison of FCCs as a scatter plot. Most FCCs are close to the diagonal in Figure 8, demonstrating that the overall control of the models is similar. Both models also exhibit differences with respect to metabolic control, in particular with respect to reactions related to ATP production and consumption. Key differences are:

**Fig. 7.**
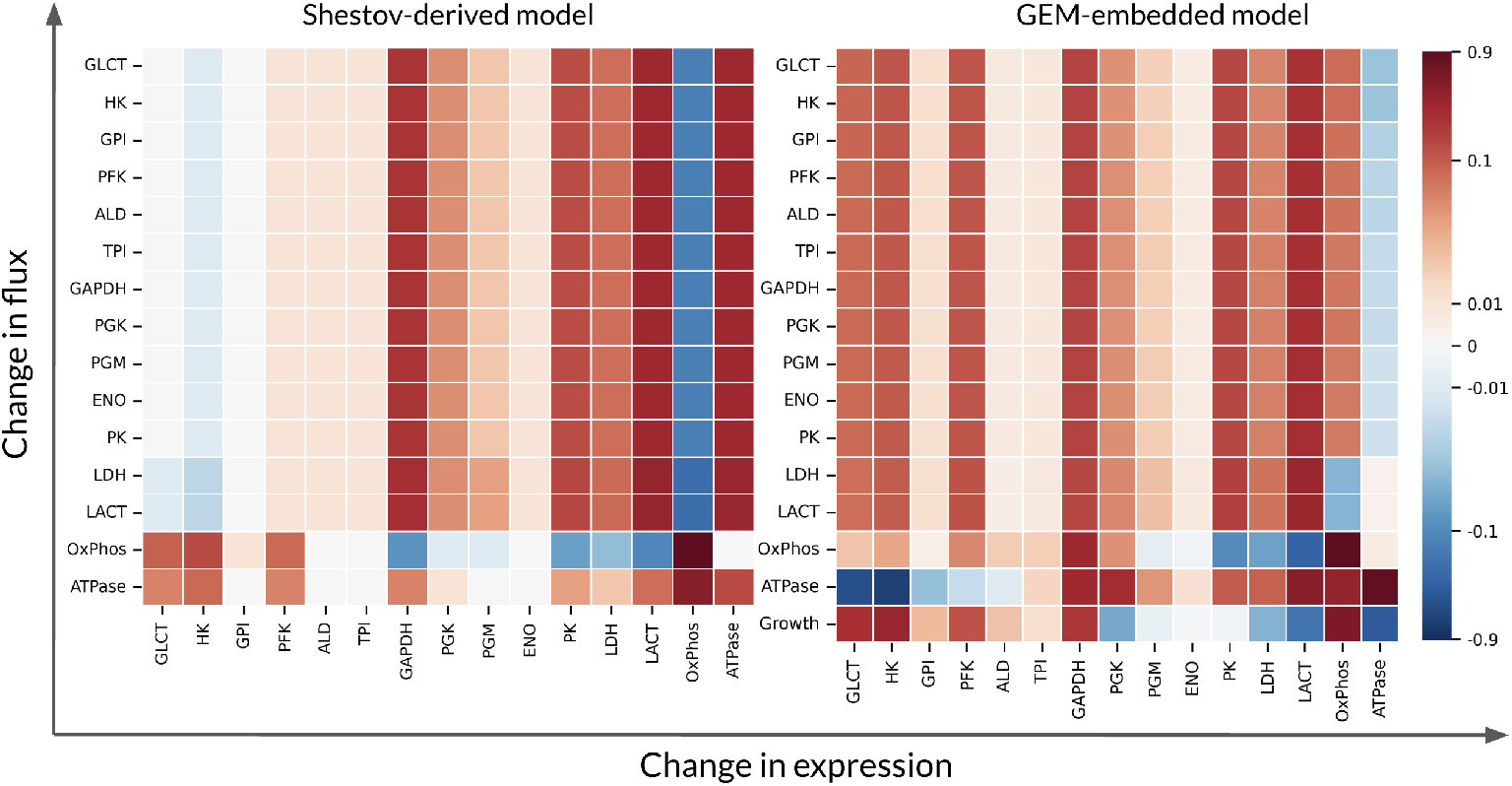
A global comparison of FCCs between the Shestov-derived model and the GEM-embedded model. Blue colors indicate negative FCCs, while red colors denote positive FCCs. Within the matrix, the x-axis corresponds to enzymes whose expression is changes, the y-axis corresponds to the reactions whose flux is affected.

**Fig. 8.**
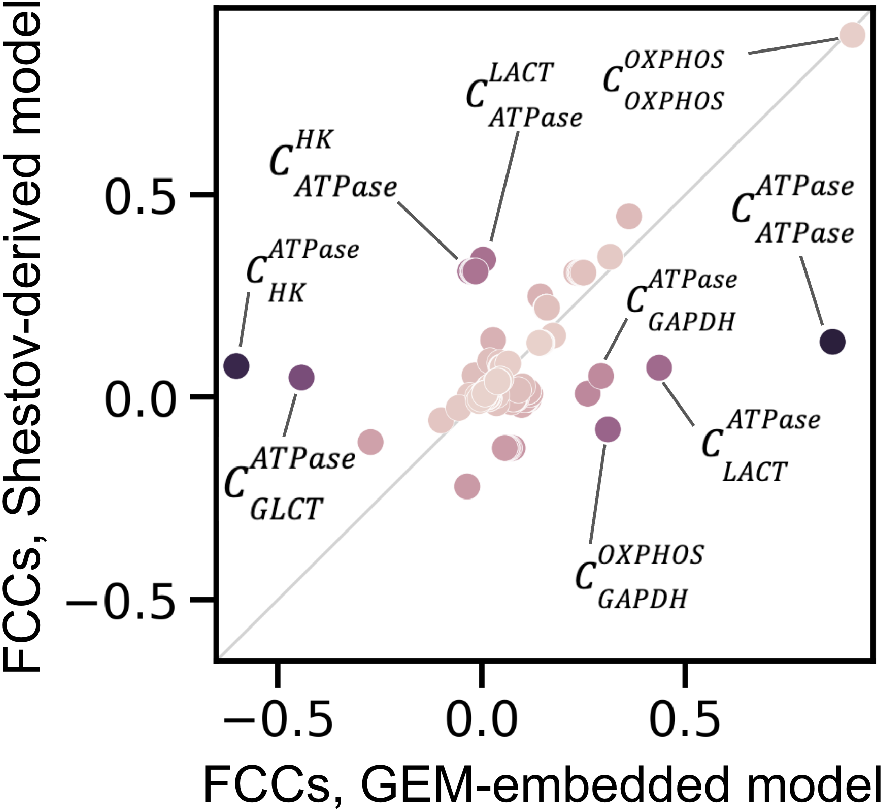
A global comparison of FCCs shown in Fig. 7 between the Shestov-derived model and the GEM-embedded model. Shown is a scatter plot of all FCCs present in both models. Most values are close to the diagonal. Dots are color-coded based on the difference in respective FCCs between the two models.

- **Control of ATPase flux:** In the model derived from Shestov et al (2014), all enzymes have a positive or low FCC on ATPase flux (bottom row in Figure 7 left side). That is, an increase of enzyme expression results in either increased ATPase flux (as the sole energy sink) or has minor impact on ATPase flux. In contrast, within the GEM-embedded model ATPase flux *decreases* upon increased expression of enzymes in upper glycolysis (GLCT, HK, GPI, PFK, and ALD). Similar to Shestov et al., however, enzymes of lower glycolysis, in particular GAPDH, have a positive FCCs on ATPase flux. Highest control on ATPase in the GEM-embedded model is the NGAM-associated reaction itself showing that flux through the ATPase reaction is largely determined by ATP demand, whereas in the Shestov-derived model, the highest control is exerted by OxPhos reaction.
- **Control of OxPhos flux:** In both models, the highest control on OxPhos is exerted by the expression of the associated enzymes itself. That is, mitochondrial flux is limited by the mitochondrial capacity, as also assumed in the parameterization. Likewise, in both models, LACT exerts negative control on OxPhos. The most pronounced difference is the control exerted by GAPDH with a positive FCC in the GEM-embedded model but a (small) negative FCC in the model derived from Shestov et al (2014).
- **Control of glycolytic flux:** Overall control is similar in both models. In the model derived from Shestov et al (2014), GAPDH exerts a strong positive control over the glycolytic flux (see the corresponding column in Figure 7), recapitulating a key result of their work (‘flux through the enzyme GAPDH is a limiting step’). Within the GEM-embedded model, GAPDH likewise exerts strong positive control, however, other reactions, such as GLCT, HK, PFK and PK exert a similar positive control. A key difference is the low control of ATPase on glycolytic flux in the GEM-embedded model, compared to the Shestov-derived model, again due to the ATPase being the sole energy sink in the latter.
- **Control of the growth reaction:** The Shestov-derived model includes no explicit representation of growth, hence its control can not be evaluated. Within the GEM-embedded model, ATPase has strong negative control on the growth reaction, since ATPase is a competing energy sink. Within the GEM-embedded model, control of the growth reaction by glycolytic enzymes is typically inverse to the control of the ATPase reaction as can be seen in a comparison of the bottom two rows in Figure 7 right side, i.e., an enzyme with positive control on the ATPase flux mostly has negative control on growth, as expected for competing reactions. Exceptions are GAPDH and OxPhos. OxPhos exerts a strong positive control on growth as well as on the ATPase. The FCC of GAPDH on growth as well as on ATPase confirms the strong impact of GAPDH on overall glycolytic flux. Different to its impact on glycolytic flux, PK exerts a very small negative control on growth, possibly because of its position in lower glycolysis such that it withdraws precursors from the growth reaction. The control of growth is further discussed below.

### Robustness and Monte-Carlo Analysis

The model derived from Shestov et al (2014), although still comparatively small, already comprises 75 kinetic parameters. Identifying these parameters remains challenging even with the increasing availability of large quantitative experimental datasets, machine learning (Li et al, 2022; Choudhury et al, 2022; Kroll et al, 2023), and dedicated databases (Chang et al, 2021; Wittig et al, 2017). To facilitate a comparison between the GEM-embedded model and the Shestov-derived model, we followed the paramerization given in Shestov et al (2014), and utilized the same set of parameters for all reactions common to both models.

In the following, we seek to evaluate the robustness of model results with respect to different choices of kinetic parameters. To this end, we make use of Monte-Carlo (MC) sampling of kinetic parameters, based on a previously established method (Steuer et al, 2006; Steuer and Junker, 2009; Murabito et al, 2014). To this end, the steady state metabolite concentrations are assigned to steady-state reference values obtained from the kinetic model analyzed in the previous section. The steady state flux distribution is obtained from the FBA solution and likewise corresponds to the metabolic state analyzed in the previous sections. All flux and concentration values are provided in Supplementary section.

For both models, we then sample the Michaelis-Menten parameters of all enzymatic reactions within two orders of magnitude around their respective original value. Each sampled set of parameters gives rise to a model instance while retaining the concenctration and flux values at the prescribed metabolic state. Each model instance is then tested for stability of the steady-state by evaluating the eigenvalues of the Jacobian matrix. For each stable model instance, the FCCs of all enzymes are evaluated.

Figure 9 shows the resulting distribution of FCCs of enzymes on the HK flux for both model ensembles, analogous to Figure 6 for the original parameterization. The medians of the distributions are in good agreement with the values obtained for the original parameterization. The distributions of FCCs of reactions close to equilibrium (i.e., GPI, ALD, TPI, PGM, ENO) are narrow and close to zero, reflecting the fact that enzymes of reactions close to equilibrium typically do not exert large flux control (Steuer and Junker, 2009; Murabito et al, 2014).

**Fig. 9.**
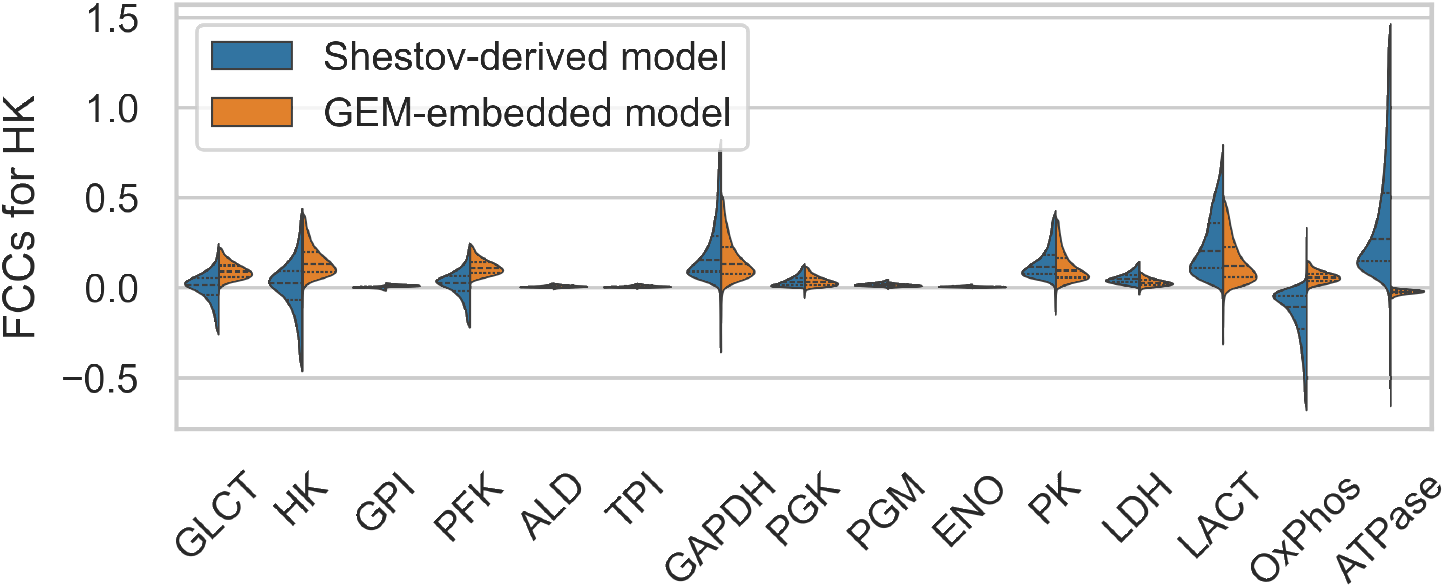
Comparison of the distribution of FCCs for HK flux between the sampled model instances of the Shestov-derived model and the sampled model instances of the GEM-embedded model.

Several enzymes exhibit a broad distribution of FCCs on HK in both model ensembles, in particular GLCT, HK, GAPDH, PK and LACT. Notably, the median of the distributions of the FCCs of OxPhos on HK exhibit opposing signs in both models, indicating that the different signs already observed in Figure 6 are a result of the model structure and the metabolic state and not the particular choice of kinetic parameters. The FCC of the ATPase on HK exhibits a wide distribution for the Shestov-derived model, indicating a high dependence on the parameterization, whereas the distribution in the GEM-embedded model is narrow and with small absolute values. The results shown in Figure 9, in comparison with Figure 6, confirm that the FCCs are mostly not strongly sensitive to the specific choice of Michaelis-Menten parameters and are constrained by network topology and the metabolic state.

Going beyond control of the HK reaction, Figure 10 provides a global view on the distribution of FCCs for the GEM-embedded model ensemble. The distributions reflect the results obtained for the original parameterization (as shown as in Figure 7, right panel). For comparison, the values of the FCCs of the original parameterization are depicted as red lines. The corresponding figure for the model derived from Shestov et al (2014) is provided as Figure S3. Small values of FCCs obtained for the original parameterization typically correspond to narrow distributions of FCCs for the model ensemble, in particular for reactions close to equilibrium (e.g., GPI, ALD, TPI, PGM, ENO) with absolute values close to zero. Large FCCs in the original parameterization correspond to broad but often sign-dominant distributions, i.e., even for broad distributions, most FCCs within the distribution have the same sign. The latter is further illustrated by the sign distribution directly (provided as Supplementary Figure S4 for the GEM-embedded model ensemble and as Figure S5 for the model ensemble corresponding to the Shestov-derived model).

**Fig. 10.**
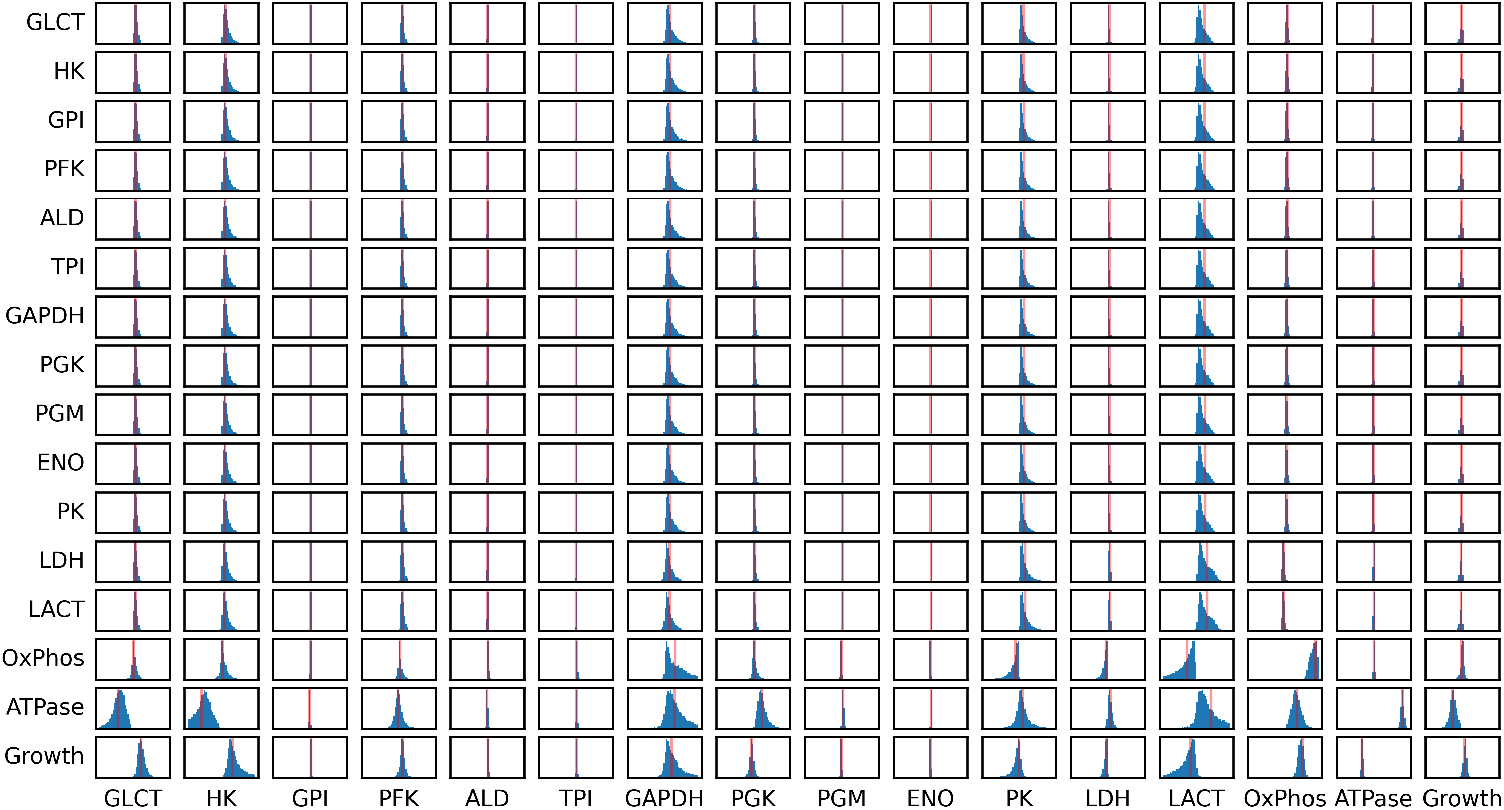
Distribution of FCCs for the GEM-embedded model. Each column shows the FCC corresponding to how a change in the expression of an enzymatic step (or coarse-grained reaction) affects the fluxes. The individual distributions of FCCs is shown in [−1, 1] intervals. The red line indicated the FCC obtained for the set of reference parameter, as shown in Figure 7B.

We emphasize that the distributions of FCCs shown in Figure 10 represent possibilities in the context of uncertain or unavailable information about enzyme-kinetic parameters, while assuming that the metabolite concentrations are constrained close to values reported by König et al (2012). A broad distribution does not imply a particular FCC is necessarily large. However, we argue that the distributions can provide a good indication about the overall structure of flux control in a metabolic model and are useful to highlight differences between models, irrespective of the choice of parameterization.

### The control of cellular growth

Of particular interest is the control of the specific growth rate in the GEM-embedded model, as pathway-focussed cancer models so far do typically not provide an explicit description of the growth rate.

Figure 11 shows the distribution of FCCs on the growth reaction for the GEM-embedded model ensemble. The figure confirms the possibility of strong control of GAPDH on growth supporting its role as a possible drug target (Shestov et al, 2014). The distribution, however, also indicates model instances of weak positive or even negative control, depending on the parameterization. Likewise, the distribution of the FCC of PK on growth indicates an ambiguous control, i.e., depending on the specific parameters the impact might be either positive or negative, with negative impact likely caused by the withdrawal of glycolytic precursors required for growth towards pyruvate.

**Fig. 11.**
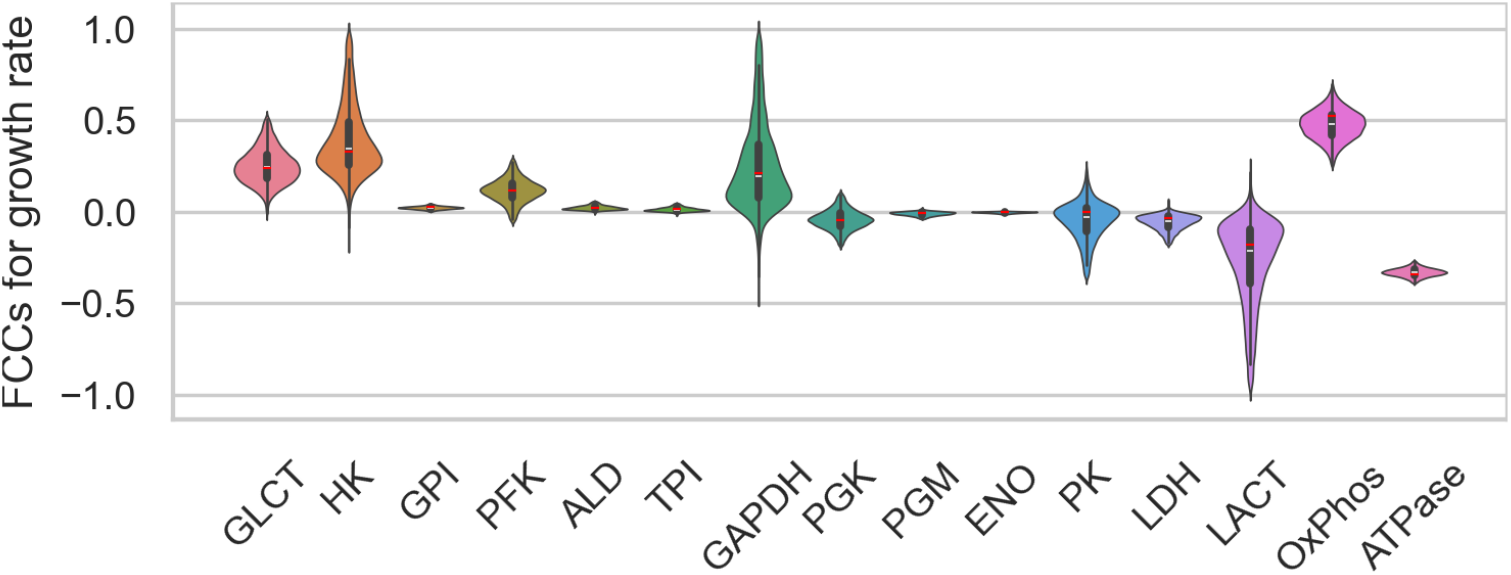
Distribution of FCCs on growth rate in the GEM-embedded model.

HK has a potentially strong control over growth, different from its control over ATPase (see Fig 10 for comparison). The ATPase itself has negative control over growth, as ATP utilization is competing with ATP use for cellular growth, but the ATPase has no strong control over the fluxes in glycolytic reactions (see Fig 10).

As expected, the OxPhos reaction exerts positive control, whereas LACT exhibits the possibilty of negative control on growth. The different signs of the FCC of the OxPhos and LACT reaction on the growth rate reflects the different yields in terms of ATP generation, as well as the competition of both reactions for pyruvate.

Overall, Figure 11 shows that the growth rate in the GEM-embedded model has a distinct control profile that does not necessarily match the control profile of key glycolytic reactions. Hence, the inclusion of cellular growth as an explicit process allows us to gain more specific insights in the control properties of the pathway.

### Conclusions

The purpose of this work was to outline a computational framework for the generation of a GEM-embedded kinetic model and to exemplify this framework by developing a kinetic model of cancer metabolism. Our aim was to overcome limitations of current pathway-focused kinetic models of metabolism and to incorporate the information provided by recent high-quality genome-scale reconstructions of human metabolism (Robinson et al, 2020). Specifically, genome-scale metabolic reconstructions provide a comprehensive and stoichiometrically correct account for the withdrawal of metabolic intermediates for growth, allowing us to embed pathway-focused models into a metabolic whole-cell context.

The proposed workflow combines several previously developed methods. Our starting point was a cell-type-specific model extracted from a current genome-scale reconstruction Agren et al (2012). The cell-type-specific model is then stoichiometrically reduced such that a set of desired metabolic functions is retained (Erdrich et al, 2015; Röhl and Bockmayr, 2017; Ataman et al, 2017). Reactions that are not part of the protected core model are aggregated into coarse-grained overall reactions, as exemplified here with the coarse-grained growth reaction and oxidative phosphorylation. We note that, while there are a number of stoichiometric reduction algorithms implemented (Singh and Lercher, 2020), they often include only the removal of reactions, without subsequent compression implemented in Erdrich et al (2015).

Parameterization of the protected core reactions follows established methods, while the parameterization of coarse-grained overall reactions incorporates biological knowledge, such as whether the respective flux is proteome- or substrate-limited. Here, we assumed that biosynthesis and the growth reaction is primarily limited by the availability of metabolic precursors (Fig. 4A), whereas mitochondrial respiration (OxPhos) was assumed to be limited by mitochondrial capacity (Fig. 4B).

The resulting GEM-embedded kinetic model was analyzed with respect to its stoichiometric and kinetic properties. A key advantage of our approach is the explicit incorporation of growth as a function within the model. Traditional kinetic models often approximate growth indirectly, typically by linking it to ATP consumption. In our model we can explicitly study the dependency between growth and the external glucose concentration or glucose transporter expression, leading to physiologically relevant predictions. We argue that this explicit representation allows for a more accurate assessment of metabolic control properties and potential drug targets.

Metabolic control analysis revealed key differences in how various enzymes regulate energy production and consumption within the GEM-embedded model compared to the reference model derived from Shestov et al (2014). Notably, while most enzymes in the reference model positively regulate ATPase flux, our model demonstrates that upper glycolytic enzymes (GLCT, HK, GPI, PFK, ALD) negatively control ATPase flux, reflecting a more distributed energy regulation. Regarding glycolytic flux, both models indicate GAPDH as a major control point. However, in particular within the GEM-embedded model, control of glycolytic flux is distributed and shared across several enzymes and reactions, such as HK, PFK, and LACT (Fig. 7).

Within the GEM-embedded model, ATPase has less control over glycolytic flux. Rather, with the explicit representation of growth, ATPase acts as a competing energy sink, exerting negative control on growth.

To account for uncertainties in kinetic parameters and to test the robustness of the results independent of the parameterization, we implemented a Monte Carlo (MC) sampling approach. This technique allowed us to assess the robustness of our results across a wide range of parameter values, confirming that our findings reflect fundamental properties of the metabolic network.

Overall, our study presents a suitable approach for embedding kinetic models within genome-scale networks, facilitating a more comprehensive and accurate representation of cancer metabolism. We have shown that explicitly modeling growth and incorporating stoichiometric constraints by incorporating genome-scale metabolic models into a kinetic GEM-embedded model can improve metabolic simulations and subsequent analysis of control properties, including the analysis of potential intervention points and drug targets.

## Materials and Methods

### Software tools

Several software tools and scripts were employed to process and analyze different types of models and data. To extract the cell-specific model from a generic GEM, we used a MATLAB implementation of tINIT algorithm (Agren et al, 2014) from the Raven Toolbox v2.6.1 (https://github.com/SysBioChalmers/RAVEN).

For the analysis of constraint-based models, we applied the COBRApy package (v0.23.0) (Ebrahim et al, 2013). The scripts for network reduction and post-processing were developed in MATLAB using the NetworkReducer implementation from the CellNetAnalyzer package (v2021.1) (von Kamp et al, 2017). To speed up the reduction process, we introduced parallelization into the CNAreduceMFNetwork routine of CellNetAnalyzer (https://gist.github.com/stairs/8dadb9f260a7354dc51db16af10e81f9). We note that the pruning step of NetworkReducer is identical to MinNW (Röhl and Bockmayr, 2017), therein formulated as a Mixed-Integer Linear Programming (MILP) problem.

For creation and analysis of kinetic models, we used COPASI v4.34 (Hoops et al, 2006). All developed kinetic and constraint-based models were uploaded to GitHub in standard SBML format (Hucka et al, 2003). The source code and data necessary to reproduce the workflow and generate the figures in this paper are available at https://github.com/stairs/GEM-reduction-for-kinetic-modeling.

### Workflow

As a starting point for developing a stoichiometrically-correct dynamic model of cancer metabolism, we selected version 1.11.0 of the Human-GEM lineage. As a comprehensive representation of the entire organism metabolism, Human-GEM includes metabolic reactions occurring in any human cell type. Since the study is focused on cancer cell metabolism, we first isolated a GEM subnetwork that would be representative of a cancer cell line. To this end, we performed GEM extraction using the expression profile of the HT29 cell line from DepMap. The data was retrieved using gdc-client tool (https://gdc.cancer.gov/) and converted to transcripts per million (TPM).

For contextualization of the model, we employed the tINIT algorithm (Agren et al, 2014). To determine which reactions from the original model should be included, we used RNA-Seq data with an expression threshold of 1 TPM. Since tINIT requires the input model to have boundary metabolites, we added reactants (or products) to all sink reactions before running the algorithm. In addition, the tINIT tool provides the ability to specify which metabolic tasks the contextualized model should be able to perform. For this purpose, we used the 57 metabolic tasks proposed by Agren et al (2014), i.e., key metabolic functions categorized as energy and redox (ER), internal conversions (IC), substrate utilization (SU) and biosynthesis of products (BS). Examples of tasks include aerobic rephosphorylation of ATP from glucose, oxidative phosphorylation, beta oxidation of long-chain fatty acids, de novo synthesis of nucleotides, uptake of essential amino acids, among others (Agren et al, 2014). For optimization, the Gurobi Optimizer version 9.0.1 was employed. The execution time of tINIT algorithm was about 50 minutes on a machine with Quad-Core Intel Core i7, 2.2 GHz.

To define the nutritional constraints, we used limits for the essential nutrients and then minimized the fluxes with the help of pFBA. The maximum uptake rates for all media components were set according to the limits chosen by Folger et al (2011), where each type of compound is assigned its average biologically justified value. Thus, for example, vitamin intake is limited to 0.005 mmol/gDW/h, amino acids to 0.05 mmol/gDW/h, while glucose consumption is allowed up to 5 mmol/gDW/h. The full list of applied medium constraints is presented in Table S1. Under the given constraints, the growth rate is limited not by glucose uptake but by other essential nutrients. The FBA problem was formulated such that we obtain the solution with minimal glucose uptake at the maximal growth rate for the given medium constraint.

### Ensuring consistent units of the growth reaction

When integrating the growth reaction from the reduced network into the kinetic model, we must ensure consistent units in the definition of the growth reaction. Constraint-based analysis typically measures fluxes in mmol/gDW/h, whereas eperimental works typically use volume (mM/h). The respective conversion factor was calculated for DB1 cells, which were instrumental in parametrisation of the model derived from Shestov et al (2014). According to Shestov et al (2014), 10^9^ DB1 cells yield around 200 mg of dry mass, and the volume of 5 × 10^8^ cells constitute approximately 1 ml. The DB1 cell dry mass and volume are therefore, on average, 0.2 ng and 2 pl, respectively. Hence, the conversion factor for the stoichiometric coefficients from mmol/gDW to mmol/l was set to 0.01 l/gDW.

The definition of the growth reaction is analogeuos to the definition of used in FBA. The mass balance of a growing cell is given as

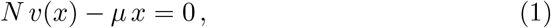

where *x* denotes the concentrations of biomass compounds and *v* is the vector of reaction fluxes (measured in mmol/gDW/h). Per definition, 1 gDW of biomass has to contain 1 g of biomass components, hence *ρ*^*T*^ *x* = 1 with *ρ* denoting the molar mass of each component. For a given vector of concentrations *x*^*\**^, the rate of biomass formation is *v*_BM_ = *µ x*^*\**^ (measured in mmol/gDW/h). within the main text, we refer to the growth rate as *µ* (measured in units 1/h) that is obtained from *v*_BM_ by multiplication with gDW/mmol.

### Metabolic Control Analysis

Metabolic control analysis (MCA) was employed to analyze how embedding of the model into a broader metabolic setting affects the distribution of flux control. Specifically, we computed the scaled or normalized flux control coefficients (Reder, 1988; Steuer and Junker, 2009), denoted as 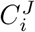,

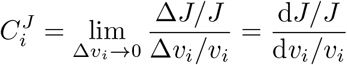

The flux control coefficients measure the ratio between the relative change of the steady-state flux *J* and the relative change in the activity of perturbed enzyme *i*, caused by an infinitesimal increase in a parameter associated with maximum enzyme activity. For the model parametrised with the reference parameter set, MCA was conducted using COPASI v4.34 (Hoops et al, 2006).

For Monte Carlo sampling, we followed the formulation derived by Reder (1988) to compute the matrix *C*^*J*^ describing the flux control coefficients

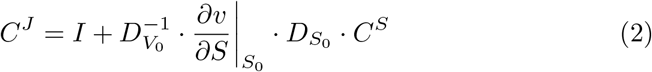

where *I* denotes the identity matrix, while 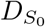 and 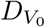 are diagonal matrices incorporating steady-state concentrations *S*_0_ and fluxes *V*_0_ along their main diagonals. The matrix *C*^*S*^, which represents concentration control coefficients, is defined as

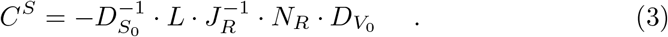

Here, *N*_*R*_ denotes the stoichiometric matrix of the reduced system, *L* is the link matrix, and *J*_*R*_ represents the Jacobian of the reduced system which is computed as

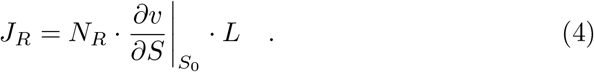

See also Steuer and Junker (2009) for definitions and details. Applying equation (2) for FCC computation only requires knowledge of the stoichiometric matrix, the flux vector *V*_0_, the vector of steady state concentrations *S*_0_, as well as the elasticities, thereby circumvents the need for numerical integration of the set of differential equations. In the main text all reported control coefficients refer to normalized (scaled) values and express relative changes.

### Metabolic control of flux ratios

MCA can be used to quantify the impact of enzyme activity on flux ratio. If we are interest how the ratio *v*_*A*_*/v*_*B*_ of two metabolic fluxes changes a function of the expression of an enzyme, the normalized control coefficient can be calculated as,

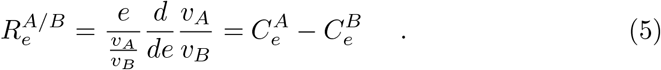

Figure S6 shows the normalized control coefficient for the ratio *v*_*LACT*_ */v*_*GLCT*_ in the GEM-embedded kinetic model.

### Monte-Carlo Sampling

The probabilistic analysis is based on the method detailed in Murabito et al (2011, 2014), following earlier approaches (Wang and Hatzimanikatis, 2006a,b; Steuer et al, 2006; Steuer and Junker, 2009). The Monte-Carlo sampling approach relies on the premise that although exact values of kinetic parameters are often not available, a metabolic state of interest, defined by steady-state concentrations *S*_0_ and fluxes *V*_0_, can often be experimentally obtained. Therefore the sampling strategy for kinetic parameters is designed to replicate the metabolic state (*S*_0_, *V*_0_) for each sampled parameter set.

In our adaptation of the method proposed by Murabito et al (2011, 2014), we start by randomly sampling each Michaelis-Menten kinetic parameter from the interval [10^*−n*^ *·* 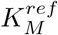, 10^+*n*^ *·* 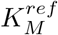], where *n* = 1 and 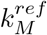 denotes the reference value of the respective parameter, as defined in the original model (Shestov et al, 2014). To ensure an even coverage across, parameter values are drawn from a logarithmic distribution.

For each reaction rate with the functional form *v* = *v*_max_ · *f* (*S, K*_*M*_), we then determine the value of *v*_max_ such that the steady state flux *V*_0_ in this reaction is preserved for each sampled set of parameter values, 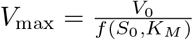.

We repeat the parameter sampling following the described routine until 10^5^ parameter sets are generated. Only parameter sets that give rise to stable models are retained. Stability of each sampled model is guaranteed by evaluating eigenvalues of the associated Jacobian matrix and by verifying that all of them have a negative real part. Following the Monte-Carlo sampling, we analyze the ensemble of sampled models within the framework of metabolic control analysis.

### Definition of the Shestov-derived model

The comparison is based on a modified version of the model of Shestov et al (2014). Compared to the original model, available on BioModels with the ID MODEL1504010000 (https://www.ebi.ac.uk/biomodels/MODEL1504010000), the model has a modified description of the respiratory OxPhos reaction. In particular, we identified a numerical instability in the original model that was caused by OxPhos and MAS (malate–aspartate shuttle) reactions sharing an identical rate law. The two reactions were then combined into a single reaction. The modification gave rise to a fourth mass conservation law, related to the oxidation state of the metabolites (see Table 1). Additionally, *v*_max,OxPhos_ was set to 5.25 *mmol/l/h* to align with the GEM-embedded model and to enable comparative analysis.

We then assign all rate equations and parameters of reactions present in both models to the rate equations and parameters used in modified model. We note that to obtain the required Michaelis-Menten constants, Shestov et al (2014) employed a heuristic approach, such that the *K*_*M*_ values of single substrate reactions were assumed to be equal to the steady-state substrate concentration of the respective metabolite, with concentrations and thermodynamic equilibrium values sourced from (König et al, 2012) and (Goldberg et al, 2004), respectively. The modified Shestov model is provided as Supplementary Data File S4 in SBML format and is referred to as ‘Shestov-derived model’ throughout the main text.

## Abbreviations and Notation

**Metabolites**

ADP: adenosine diphosphate
AMP: adenosine monophosphate
ATP: adenosine triphosphate
BPG: 1,3-bisphosphoglycerate
DHAP: dihydroxyacetone phosphate
F6P: fructose-6-phosphate
FBP: fructose-1,6-bisphosphate
GAP: glyceraldehyde-3-phosphate
GLC: glucose
G6P: glucose-6-phosphate
LAC: lactate
NAD: nicotinamide adenine dinucleotide
NADH: nicotinamide adenine dinucleotide (reduced)
O2: oxygen
PEP: phosphoenolpyruvate
PG2: 2-phosphoglycerate
PG3: 3-phosphoglycerate
Pi: inorganic phosphate
PYR: pyruvate

**Reactions**

AK: adenylate kinase
ALD: aldolase
ATPase: ATP utilization/maintenance
ENO: enolase
GAPDH: glyceraldehyde-3-phosphate dehydrogenase
GLCT: glucose transport
GPI: glucose-6-phosphate isomerase
HK: hexokinase
LACT: lactate transport
LDH: lactate dehydrogenase
OxPhos: oxidative phosphorylation
OXYT: oxygen transport
PFK: phosphofructokinase
PGK: phosphoglycerate kinase
PGM: phosphoglycerate mutase
PK: pyruvate kinase
TPI: triosephosphate isomerase

## Supplementary Text S1

**Putting models in context: deriving a kinetic model of cancer metabolism that includes cellular growth using genome-scale stoichiometric reduction**

*Komkova et al. (2025)*

## 1 List of Supplementary Files

- **Supplementary Text S1**. A PDF file with additional figures and tables.
- **Data File S1**. SBML file with the stoichiometric model of the reduced cell type-specific HT-29 cell GEM (pruned version).
- **Data File S2**. SBML file of the stoichiometric model of the cell type-specific HT-29 GEM (pruned and compressed version).
- **Data File S3**. The GEM-embedded kinetic model in SBML format.
- **Data File S4**. The kinetic model of Shestov et al (2014) used as reference in SBML format.
- **Data File S5**. A zip-archive containing the source code and data necessary to reproduce the workflow and generate the figures presented in the paper.

## 2 Medium composition for FBA and model reduction

**Table S1.** Uptake constraints of the growth medium used for the FBA simulations. Uptake constraints are according the limits given by Folger et al (2011).

| Metabolite | Maximum Uptake (mmol/gDW/h) |
| --- | --- |
| D-Glucose | 5.0 |
| L-Glutamine | 0.5 |
| O<sub>2</sub> | 1000 |
| Phosphate | 1000 |
| Fe<sup>2+</sup> | 1000 |
| Thiamin | 0.005 |
| Riboflavin | 0.005 |
| Pyridoxine | 0.005 |
| Nicotinamide | 0.005 |
| myo-Inositol | 0.005 |
| Folate | 0.005 |
| (R)-Pantothenate | 0.005 |
| Choline | 0.005 |
| Alpha-tocopherol | 0.005 |
| Biotin | 0.005 |
| Linoleate | 0.005 |
| Linolenate | 0.005 |
| L-Alanine | 0.05 |
| L-Arginine | 0.05 |
| L-Asparagine | 0.05 |
| L-Aspartate | 0.05 |
| L-Cystine | 0.05 |
| L-Glutamate | 0.05 |
| L-Histidine | 0.05 |
| L-Isoleucine | 0.05 |
| L-Leucine | 0.05 |
| L-Lysine | 0.05 |
| L-Methionine | 0.05 |
| L-Phenylalanine | 0.05 |
| L-cysteine/glyciner | 0.05 |
| L-Serine | 0.05 |
| L-Threonine | 0.05 |
| L-Tryptophan | 0.05 |
| L-Tyrosine | 0.05 |
| L-Valine | 0.05 |

## 3 GEM-embedded kinetic model: definitions

**Table S2.** List of reactions of the GEM-embedded model and (net) flux values for the reference state. The steady state growth rate for the reference parameters and external glucose [GLCex] = 5.0 mmol*/*l is 0.012 h^−1^.

| Reaction ID | Chemical equation | Flux<br>(mmol/l/h) |
| --- | --- | --- |
| GLCT | $\text{GLC}_{\text{ex}} \leftrightarrow \text{GLC}$ | 27.86 |
| HK | $\text{GLC} + \text{ATP} \rightarrow \text{G6P} + \text{ADP} + \text{H}$ | 27.86 |
| GPI | $\text{G6P} \leftrightarrow \text{F6P}$ | 27.37 |
| PFK | $\text{F6P} + \text{ATP} \rightarrow \text{FBP} + \text{ADP} + \text{H}$ | 27.17 |
| ALD | $\text{FBP} \leftrightarrow \text{DHAP} + \text{GAP}$ | 27.17 |
| TPI | $\text{DHAP} \leftrightarrow \text{GAP}$ | 26.94 |
| GAPDH | $\text{GAP} + \text{NAD} + \text{Pi} \leftrightarrow \text{BPG} + \text{NADH} + \text{H}$ | 54.02 |
| PGK | $\text{BPG} + \text{ADP} \leftrightarrow \text{PG3} + \text{ATP}$ | 54.02 |
| PGM | $\text{PG3} \leftrightarrow \text{PG2}$ | 53.45 |
| ENO | $\text{PG2} \leftrightarrow \text{PEP}$ | 53.45 |
| PK | $\text{PEP} + \text{ADP} \rightarrow \text{PYR} + \text{ATP}$ | 53.45 |
| LDH | $\text{PYR} + \text{NADH} \leftrightarrow \text{LAC} + \text{NAD}$ | 47.05 |
| LACT | $\text{LAC} \leftrightarrow \text{LAC}_{\text{ex}}$ | 47.05 |
| ATPase | $\text{ATP} \rightarrow \text{ADP} + \text{Pi} + \text{H}$ | 29.46 |
| AK | $\text{ATP} + \text{AMP} \leftrightarrow 2 \text{ADP}$ | 0.00 |
| OXYT | $\text{O2}_{\text{ex}} \leftrightarrow \text{O2}$ | 11.13 |
| OxPhos | $\text{PYR} + 14.75 \text{ ADP} + 14.75 \text{ Pi} + 3 \text{ O2} + \text{NADH} \rightarrow$<br>$\text{NAD} + 14.75 \text{ ATP}$ | 3.52 |
| Growth | $0.0007 \text{ BPG} + 0.4801 \text{ PG3} + 0.1953 \text{ DHAP} + 0.0043$<br>$\text{FBP} + 0.4712 \text{ O2} + 0.1647 \text{ F6P} + 0.07 \text{ GAP} +$<br>$0.4129 \text{ G6P} + 2.8958 \text{ NADH} + 2.415 \text{ PYR} + 62.9543$<br>$\text{ATP} \rightarrow 1 \text{ gDW Biomass} + 62.9543 \text{ ADP} + 2.8958$<br>$\text{NAD} + 64.287 \text{ Pi}$ | $0.012 \text{ h}^{-1}$ |

**Table S3.** The steady state concentrations of intracellular metabolites for the GEM-embedded model at the reference state.

| Metabolite | Concentration (mmol/l) |
| --- | --- |
| GLC | 2.3633 |
| G6P | 0.1225 |
| F6P | 0.0349 |
| FBP | 0.3925 |
| DHAP | 0.0200 |
| GAP | 0.0008 |
| BPG | 0.0153 |
| PG3 | 0.7572 |
| PG2 | 0.0430 |
| PEP | 0.2321 |
| PYR | 0.4162 |
| LAC | 15.0336 |
| ATP | 2.9217 |
| ADP | 0.0935 |
| AMP | 0.0030 |
| Pi | 5.5002 |
| NADH | 0.0032 |
| NAD | 0.5465 |
| O2 | 0.0671 |

### 3.1 GEM-embedded kinetic model including growth

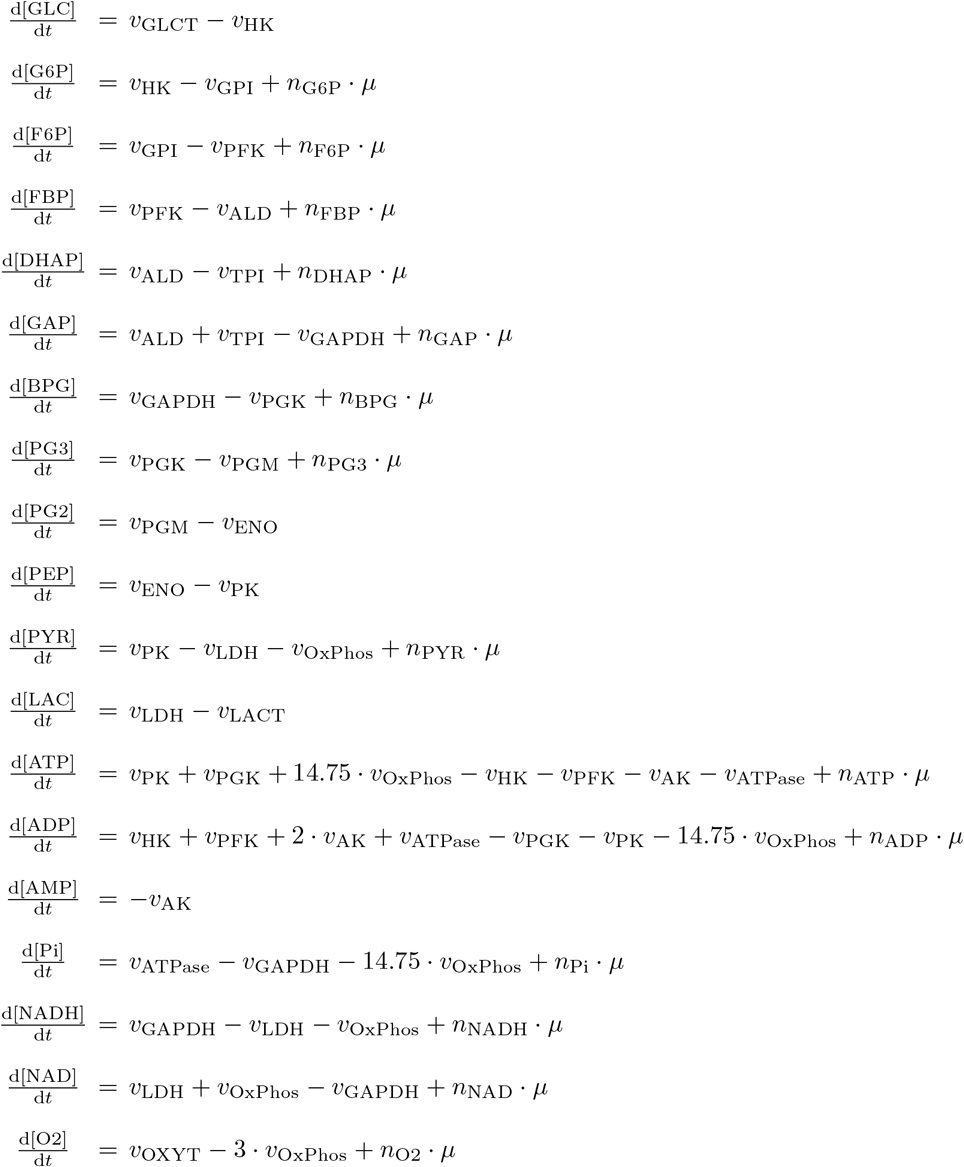

**Table S4.** Stoichiometry of the growth reaction (measured in mmol/l). We note that the values in the main text are given relative to 1 gDW. The conversion factor for the stoichiometric coefficients from mmol/gDW to mmol/l is 0.01 l/gDW.

| Stoichiometric coefficient | Value (mmol/l) |
| --- | --- |
| $n_{\text{G6P}}$ | -41.29 |
| $n_{\text{F6P}}$ | -16.47 |
| $n_{\text{FBP}}$ | -0.43 |
| $n_{\text{DHAP}}$ | -19.53 |
| $n_{\text{GAP}}$ | -7.0 |
| $n_{\text{BPG}}$ | -0.07 |
| $n_{\text{PG3}}$ | -48.01 |
| $n_{\text{PYR}}$ | -241.5 |
| $n_{\text{ATP}}$ | -6295.43 |
| $n_{\text{O2}}$ | -47.12 |
| $n_{\text{NADH}}$ | -289.58 |
| $n_{\text{Pi}}$ | 6428.7 |
| $n_{\text{ADP}}$ | 6295.43 |
| $n_{\text{NAD}}$ | 289.58 |

### 3.2 Conservation relationships

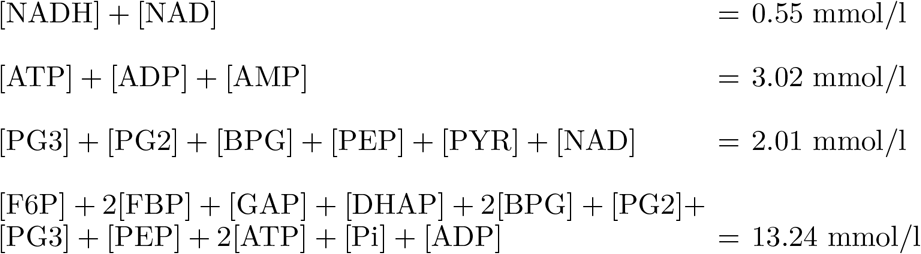

### 3.3 Reactions, rate equations, and kinetic parameters

#### Glucose transport

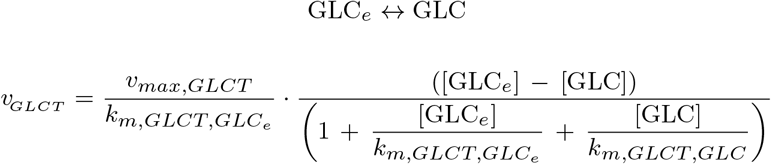

| Parameter | Value | Unit |
| --- | --- | --- |
| $v_{max, GLCT}$ | 100 | mmol/l/h |
| $k_{m, GLCT, GLC_e}$ | 2.1 | mmol/l |
| $k_{m, GLCT, GLC}$ | 2.1 | mmol/l |
| $[\text{GLC}_e]$ | 5 | mmol/l |

#### Hexokinase reaction

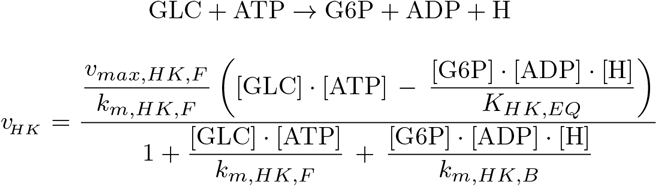

| Parameter | Value | Unit |
| --- | --- | --- |
| $v_{max, HK, F}$ | 176 | mmol/l/h |
| $k_{m, HK, F}$ | 7.5 | mmol<sup>2</sup>/l<sup>2</sup> |
| $k_{m, HK, B}$ | 2.94e-10 | mmol<sup>3</sup>/l<sup>3</sup> |
| $K_{HK, EQ}$ | 2329.1603 | mmol/l |
| $[\text{H}]$ | 1e-7 | mmol/l |

#### Glucose-6-phosphate isomerase reaction

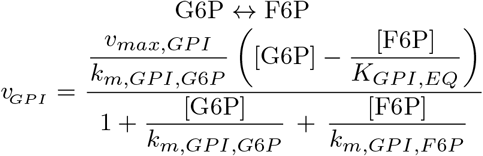

| Parameter | Value | Unit |
| --- | --- | --- |
| $v_{max, GPI}$ | 858 | mmol/l/h |
| $K_{GPI, EQ}$ | 0.3258 | - |
| $k_{m, GPI, G6P}$ | 0.25 | mmol/l |
| $k_{m, GPI, F6P}$ | 0.0773 | mmol/l |

#### Phosphofructokinase reaction

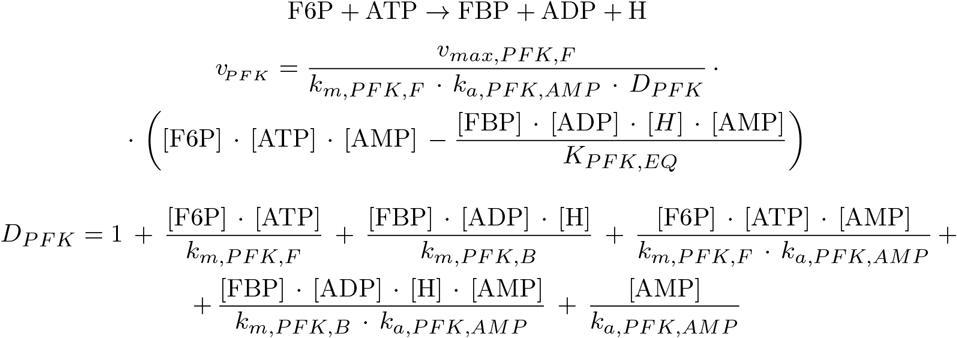

| Parameter | Value | Unit |
| --- | --- | --- |
| $v_{max,PFK,F}$ | 1769 | mmol/l/h |
| $K_{PFK,EQ}$ | 545.14 | mmol/l |
| $k_{m,PFK,F}$ | 0.23 | mmol<sup>2</sup>/l<sup>2</sup> |
| $k_{m,PFK,B}$ | 1.82e-10 | mmol<sup>3</sup>/l<sup>3</sup> |
| $k_{a,PFK,AMP}$ | 0.001 | mmol/l |
| [H] | 1e-7 | mmol/l |

#### Aldolase reaction

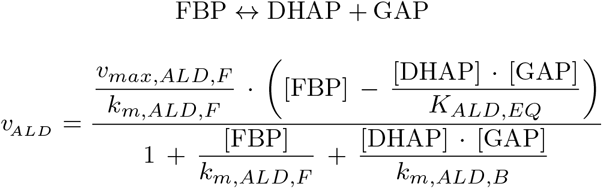

| Parameter | Value | Unit |
| --- | --- | --- |
| $v_{max,ALD,F}$ | 321 | mmol/l/h |
| $K_{ALD,EQ}$ | 4.82e-5 | mmol/l |
| $k_{m,ALD,F}$ | 0.16 | mmol/l |
| $k_{m,ALD,B}$ | 8.29e-5 | mmol<sup>2</sup>/l<sup>2</sup> |

#### Triosephosphate isomerase reaction

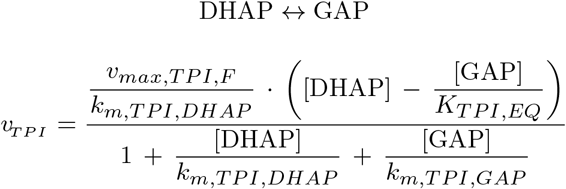

| Parameter | Value | Unit |
| --- | --- | --- |
| $v_{max,TPI,F}$ | 859 | mmol/l/h |
| $K_{TPI,EQ}$ | 0.0472 | - |
| $k_{m,TPI,DHAP}$ | 0.04 | mmol/l |
| $k_{m,TPI,GAP}$ | 0.002 | mmol/l |

#### Glyceraldehyde 3-phosphate dehydrogenase reaction

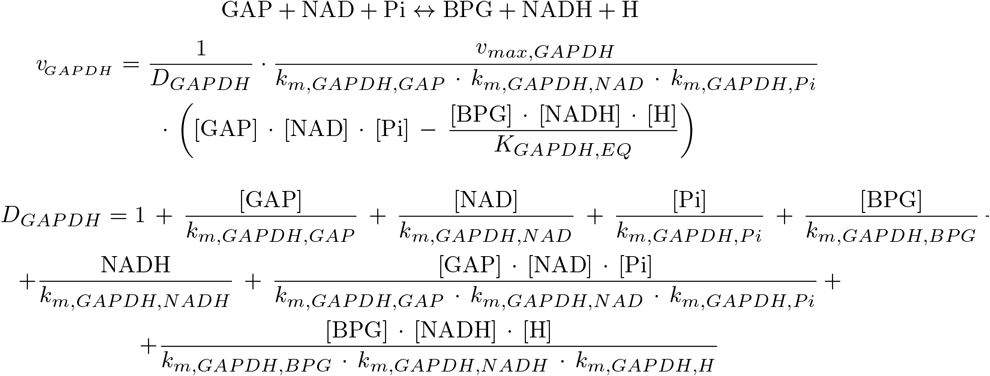

| Parameter | Value | Unit |
| --- | --- | --- |
| $v_{max,GAPDH,F}$ | 781 | mmol/l/h |
| $K_{GAPDH,EQ}$ | 0.03503 | - |
| $k_{m,GAPDH,GAP}$ | 0.002 | mmol/l |
| $k_{m,GAPDH,NAD}$ | 0.55 | mmol/l |
| $k_{m,GAPDH,Pi}$ | 4 | mmol/l |
| $k_{m,GAPDH,BPG}$ | 0.1 | mmol/l |
| $k_{m,GAPDH,NADH}$ | 0.001 | mmol/l |
| $k_{m,GAPDH,H}$ | 1e-7 | mmol/l |
| $[\text{H}]$ | 1e-7 | mmol/l |

#### Phosphoglycerate kinase reaction

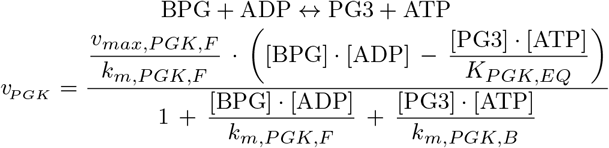

| Parameter | Value | Unit |
| --- | --- | --- |
| $v_{max,PGK,F}$ | 221 | mmol/l/h |
| $K_{PGK,EQ}$ | 6083.56 | - |
| $k_{m,PGK,F}$ | 0.00118 | mmol<sup>2</sup>/l<sup>2</sup> |
| $k_{m,PGK,B}$ | 1.5 | mmol<sup>2</sup>/l<sup>2</sup> |

#### Phosphoglycerate mutase reaction

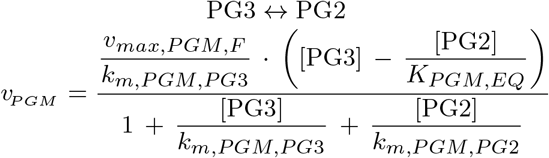

| Parameter | Value | Unit |
| --- | --- | --- |
| $v_{max,PGM,F}$ | 528 | mmol/l/h |
| $K_{PGM,EQ}$ | 0.0772 | - |
| $k_{m,PGM,PG3}$ | 0.5 | mmol/l |
| $k_{m,PGM,PG2}$ | 0.03 | mmol/l |

#### Enolase reaction

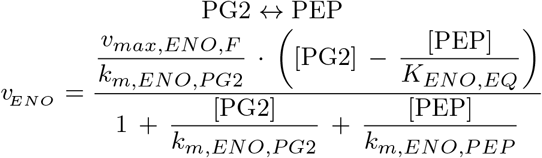

| Parameter | Value | Unit |
| --- | --- | --- |
| $v_{max,ENO,F}$ | 1340 | mmol/l/h |
| $K_{ENO,EQ}$ | 6.0689 | - |
| $k_{m,ENO,PG2}$ | 0.03 | mmol/l |
| $k_{m,ENO,PEP}$ | 0.15 | mmol/l |

#### Pyruvate kinase reaction

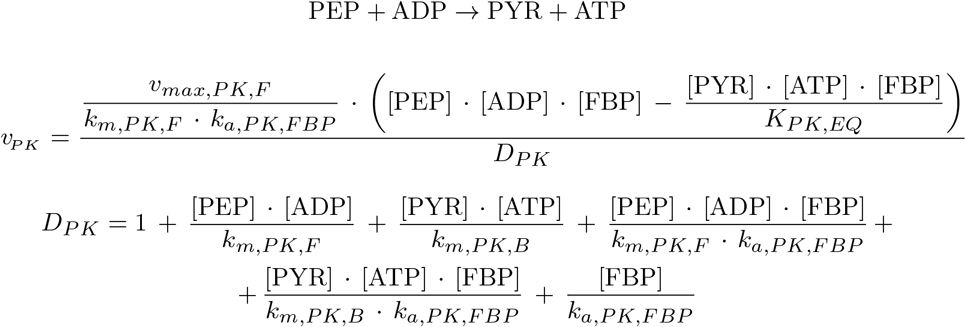

| Parameter | Value | Unit |
| --- | --- | --- |
| $v_{max,PK,F}$ | 212 | mmol/l/h |
| $K_{PK,EQ}$ | 57783.82 | mmol/l |
| $k_{m,PK,F}$ | 0.00177 | mmol<sup>2</sup>/l<sup>2</sup> |
| $k_{m,PK,B}$ | 0.15 | mmol<sup>2</sup>/l<sup>2</sup> |
| $k_{a,PK,FBP}$ | 0.5 | mmol/l |

#### Lactate dehydrogenase reaction

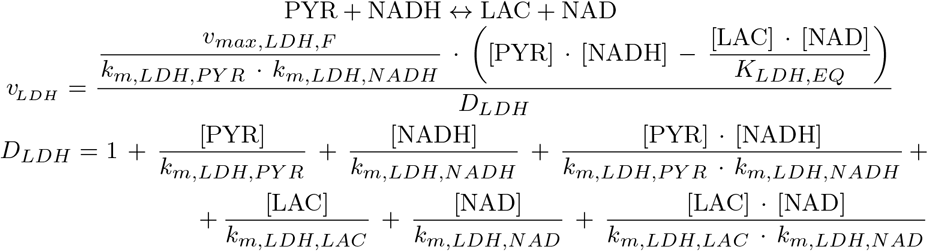

| Parameter | Value | Unit |
| --- | --- | --- |
| $v_{max,LDH,F}$ | 434 | mmol/l/h |
| $K_{LDH,EQ}$ | 15285.09 | - |
| $k_{m,LDH,PYR}$ | 0.5 | mmol/l |
| $k_{m,LDH,NADH}$ | 0.001 | mmol/l |
| $k_{m,LDH,LAC}$ | 5 | mmol/l |
| $k_{m,LDH,NAD}$ | 0.549 | mmol/l |

#### Lactate transport

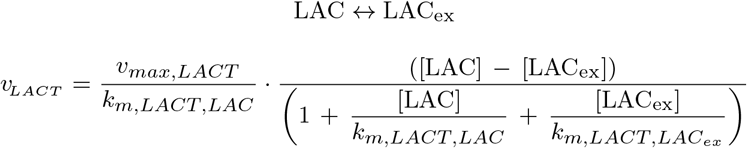

| Parameter | Value | Unit |
| --- | --- | --- |
| $v_{max,LACT,F}$ | 60 | mmol/l/h |
| $k_{m,LACT,LAC_{ex}}$ | 3 | mmol/l |
| $k_{m,LACT,LAC}$ | 3 | mmol/l |
| $[\text{LAC}_{ex}]$ | 0.5 | mmol/l |

#### ATPase reaction

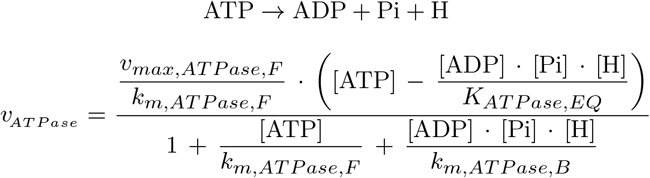

| Parameter | Value | Unit |
| --- | --- | --- |
| $v_{max,ATPase,F}$ | 390 | mmol/l/h |
| $K_{ATPase,EQ}$ | 478343.7 | mmol<sup>2</sup>/l<sup>2</sup> |
| $k_{m,ATPase,F}$ | 3 | mmol/l |
| $k_{m,ATPase,B}$ | 4.71e-9 | mmol<sup>3</sup>/l<sup>3</sup> |
| [H] | 1e-7 | mmol/l |

#### Adenylate kinase reaction

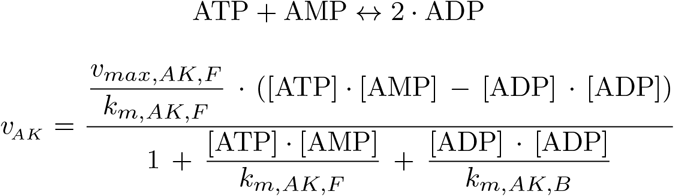

| Parameter | Value | Unit |
| --- | --- | --- |
| $v_{max,AK,F}$ | 2000 | mmol/l/h |
| $k_{m,AK,F}$ | 5 | mmol<sup>2</sup>/l<sup>2</sup> |
| $k_{m,AK,B}$ | 2 | mmol<sup>2</sup>/l<sup>2</sup> |

#### Oxygen transport

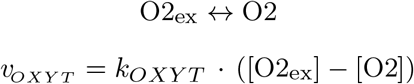

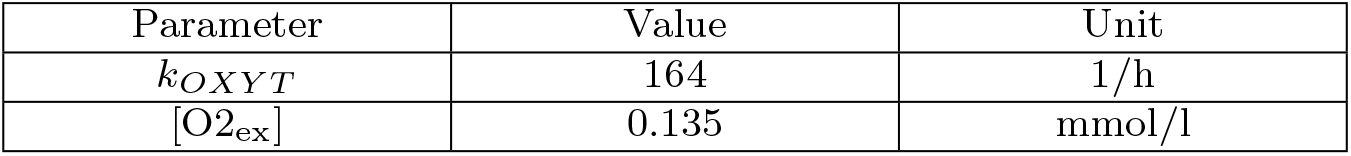

#### OxPhos reaction

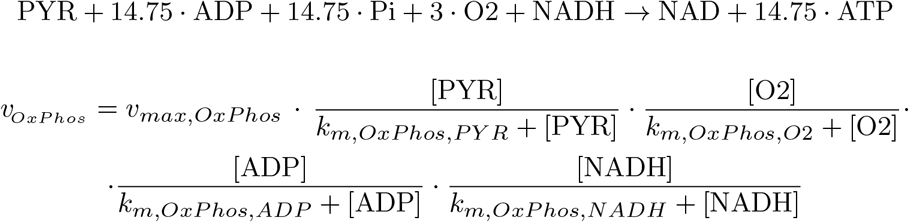

| Parameter | Value | Unit |
| --- | --- | --- |
| $v_{max,OxPhos}$ | 5.25 | mmol/l/h |
| $k_{m,OxPhos,PYR}$ | 0.001 | mmol/l |
| $k_{m,OxPhos,O2}$ | 0.005 | mmol/l |
| $k_{m,OxPhos,ADP}$ | 0.005 | mmol/l |
| $k_{m,OxPhos,NADH}$ | 0.001 | mmol/l |

#### Growth reaction

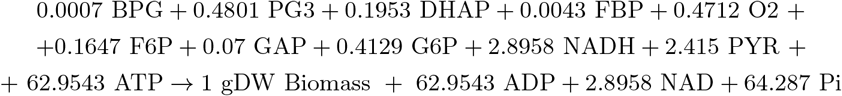

The stoichiometric coefficients of the growth reaction are measured in mmol/gDW.

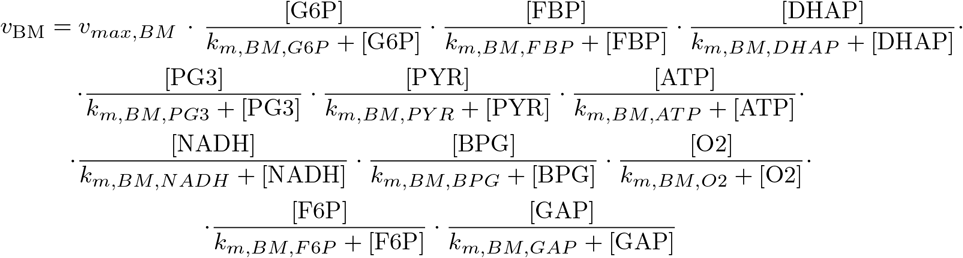

| Parameter | Value | Unit |
| --- | --- | --- |
| $v_{max,BM}$ | 200 | 1/h |
| $k_{m,BM,G6P}$ | 0.25 | mmol/l |
| $k_{m,BM,FBP}$ | 0.155 | mmol/l |
| $k_{m,BM,DHAP}$ | 0.0414 | mmol/l |
| $k_{m,BM,PG3}$ | 0.5 | mmol/l |
| $k_{m,BM,PYR}$ | 0.5 | mmol/l |
| $k_{m,BM,ATP}$ | 3 | mmol/l |
| $k_{m,BM,NADH}$ | 0.001 | mmol/l |
| $k_{m,BM,BPG}$ | 0.1 | mmol/l |
| $k_{m,BM,O_2}$ | 0.04 | mmol/l |
| $k_{m,BM,F6P}$ | 0.0773 | mmol/l |
| $k_{m,BM,GAP}$ | 0.002 | mmol/l |

## 4 Additional Figures

**Fig. S1.**
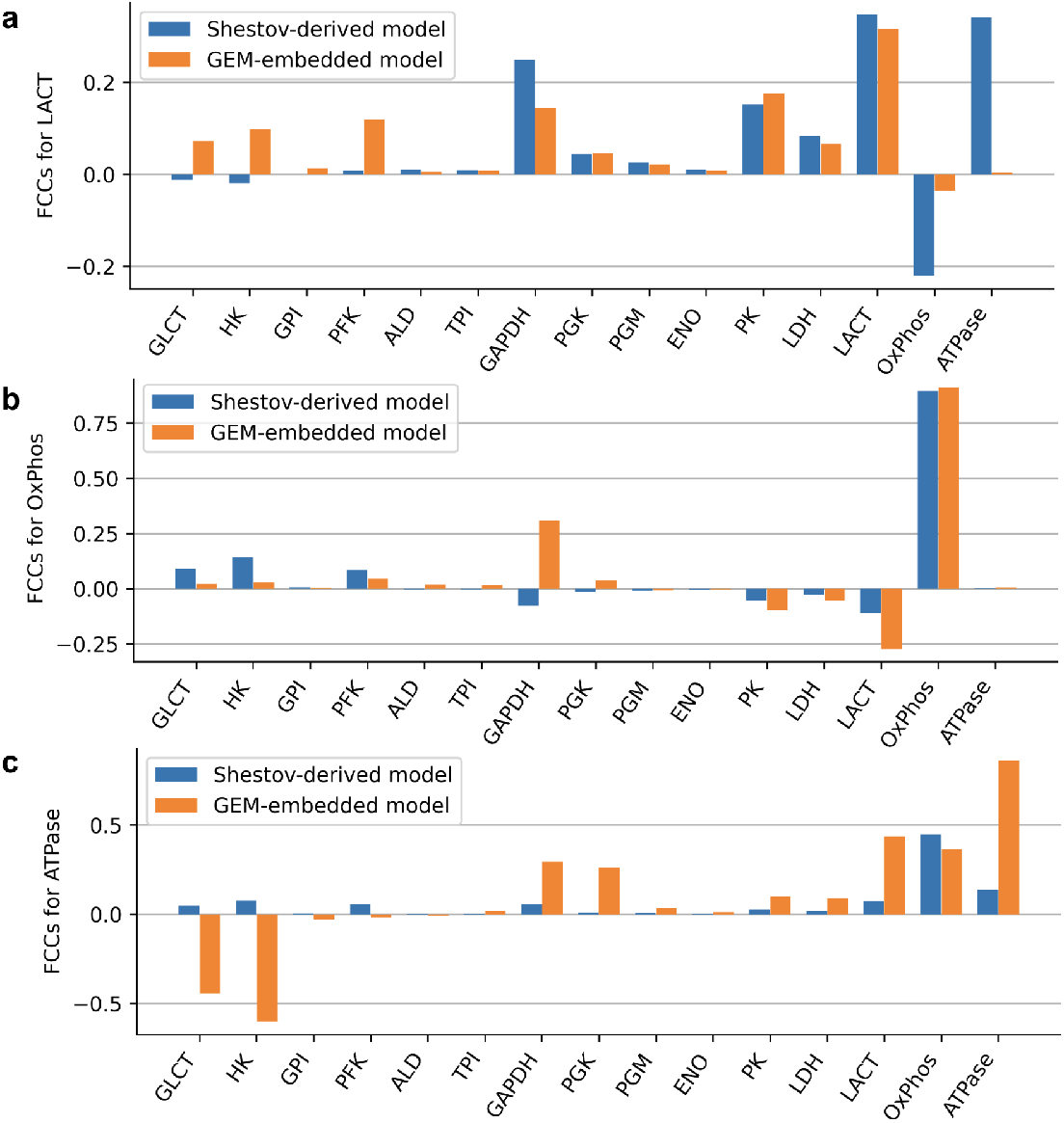
Differences in the flux control coefficients between the Shestov-derived model and the GEM-embedded model. Shown are results for the control of (A) LACT, (B) OxPhos, and (C) ATPase flux, in addition to the control of HK flux (shown in the main text).

**Fig. S2.**
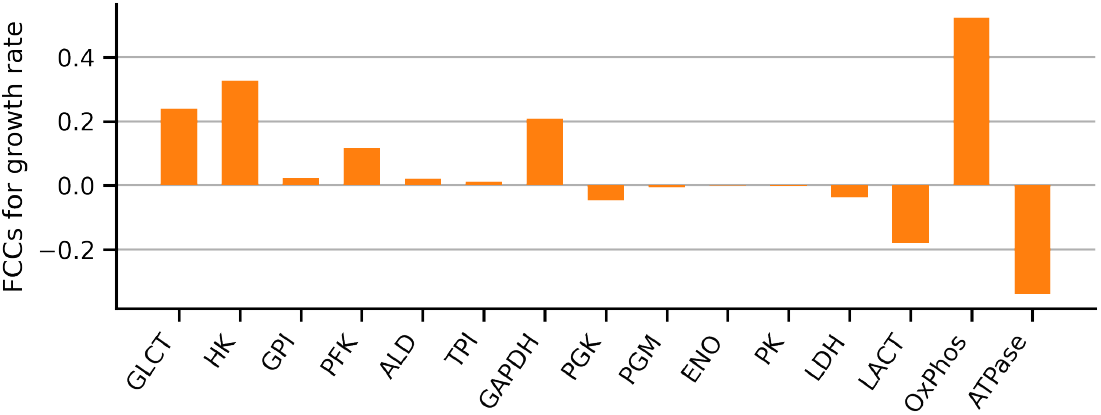
Flux control coefficients of growth rate in the GEM-embedded model.

**Fig. S3.**
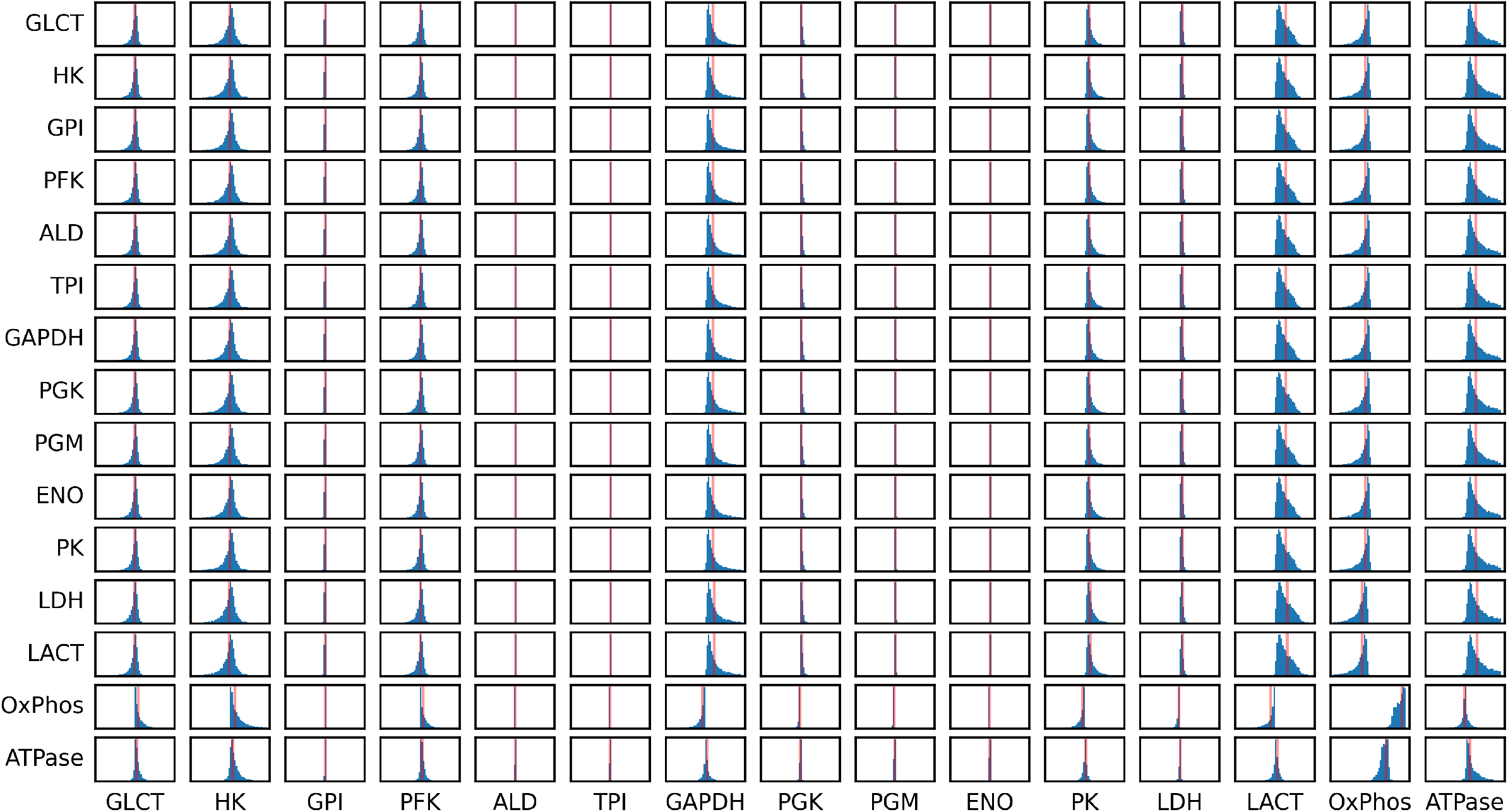
Distribution of FCCs using MC sampling of kinetic parameters in the Shestov-derived model. Columns stand for the pertubation of enzyme levels, rows represent the response in the respective flux values. The red lines show the FCCs of the model parametrized with the reference parameter set.

**Fig. S4.**
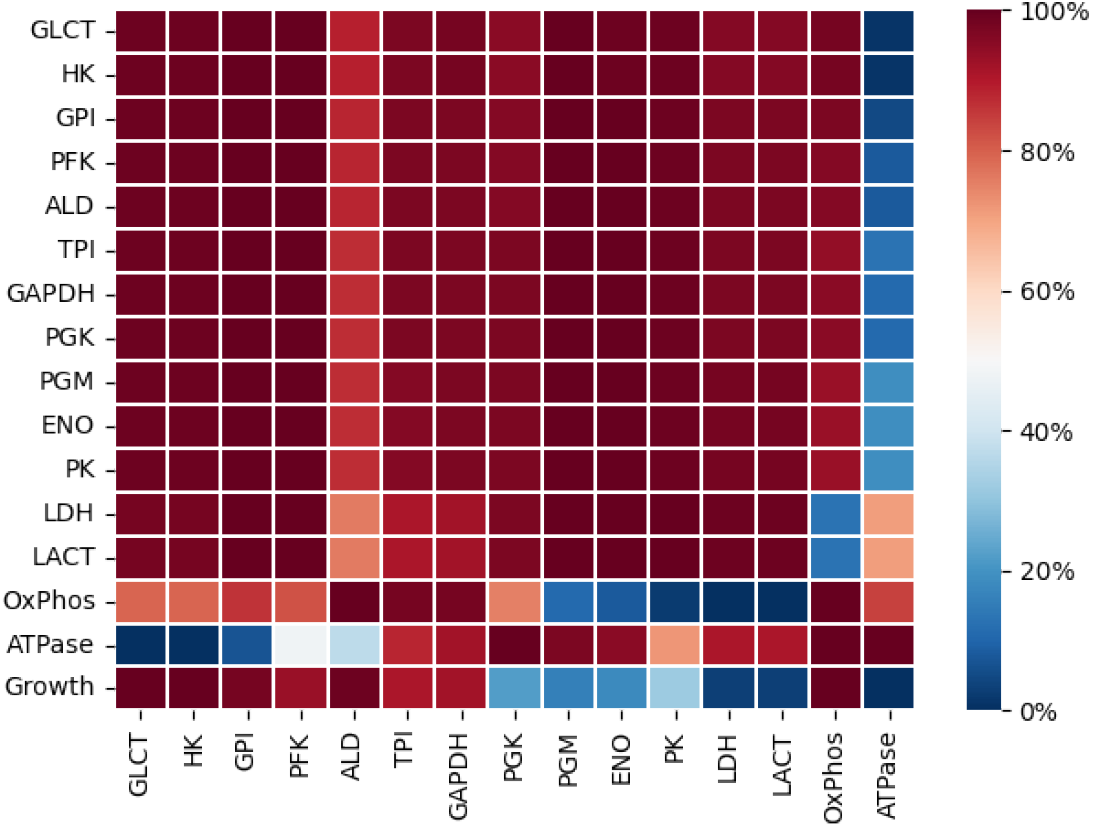
Probabilistic sign distribution of FCCs in the ensemble of GEM-embedded models. The color of the entry in the i-th row and j-th column corresponds to the percentage of FCCs with a positive values, indicating that an increase in enzyme j will result in an increase of the flux through reaction i. Most FCCs are sign-dominant, i.e., percentages are either close to 100% or close to 0%.

**Fig. S5.**
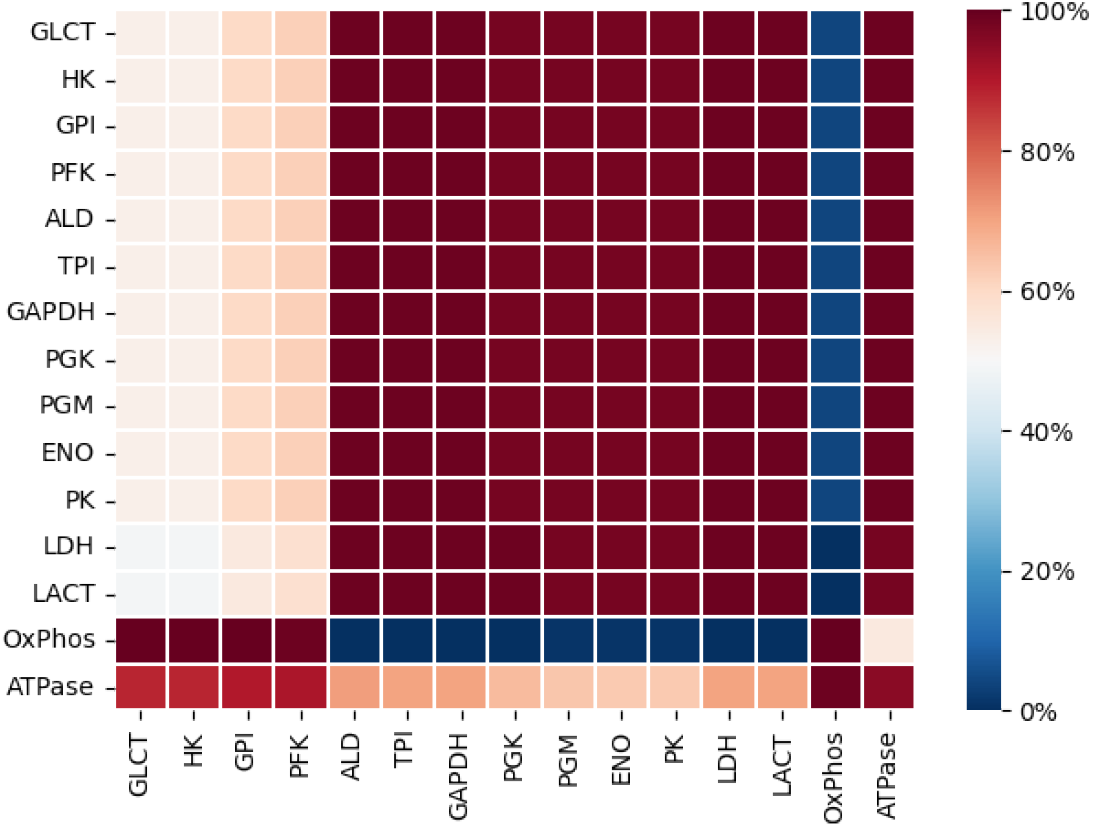
Probabilistic sign distribution of FCCs in the Shestov-derived model, corresponding to the distributions shown in Figure S3. The color of the entry in the i-th row and j-th column corresponds to the percentage of FCCs with a positive values, indicating that an increase in enzyme j will result in an increase of the flux through reaction i. Most FCCs are sign-dominant, i.e., percentages are either close to 100% or close to 0%.

**Fig. S6.**
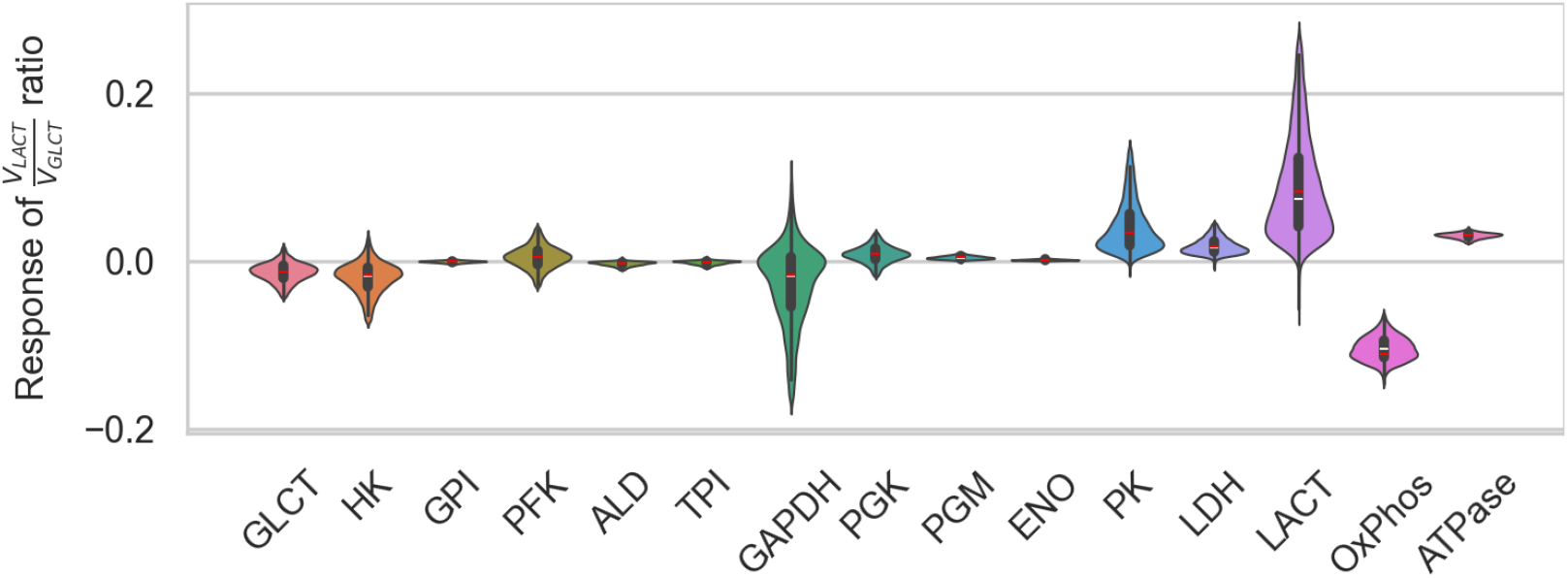
Control coefficients for the ratio *v*_*LACT*_ */v*_*GLCT*_ in the GEM-embedded model, describing the sensitivity of the ratio to a perturbation in enzyme activity. An increase in LACT will increase the ratio, whereas an increase in OxPhos (mitochondrial capacity) will decrease the ratio.

## References

Agren R, Bordel S, Mardinoglu A, et al (2012) Reconstruction of genome-scale active metabolic networks for 69 human cell types and 16 cancer types using INIT. PLoS computational biology 8(5):e1002.518. 10.1371/journal.pcbi.1002518, [Online; accessed 2024-08-09]

Agren R, Mardinoglu A, Asplund A, et al (2014) Identification of anticancer drugs for hepatocellular carcinoma through personalized genome-scale metabolic modeling. Mol Syst Biol 10:721

Ataman M, Hernandez Gardiol DF, Fengos G, et al (2017) redGEM: Systematic reduction and analysis of genome-scale metabolic reconstructions for development of consistent core metabolic models. PLoS Comput Biol 13(7):e1005.444

Baroukh C, oz Tamayo R, Steyer JP, et al (2014) DRUM: a new framework for metabolic modeling under non-balanced growth. Application to the carbon metabolism of unicellular microalgae. PLoS One 9(8):e104.499

Burns J, Cornish-Bowden A, Groen A, et al (1985) Control analysis of metabolic systems. Trends in Biochemical Sciences 10(1):16. 10.1016/0968-0004(85)90008-8, URL https://www.sciencedirect.com/science/article/pii/0968000485900088

Chang A, Jeske L, Ulbrich S, et al (2021) BRENDA, the ELIXIR core data resource in 2021: new developments and updates. Nucleic Acids Res 49(D1):D498–D508

Choudhury S, Moret M, Salvy P, et al (2022) Reconstructing kinetic models for dynamical studies of metabolism using generative adversarial networks. Nature Machine Intelligence 4(8):710–719

DeBerardinis RJ, Chandel NS (2016) Fundamentals of cancer metabolism. Science Advances 2(5). 10.1126/sciadv.1600200, URL http://dx.doi.org/10.1126/sciadv.1600200

Duarte NC, Becker SA, Jamshidi N, et al (2007) Global reconstruction of the human metabolic network based on genomic and bibliomic data. Proceedings of the National Academy of Sciences 104(6):1777–1782. 10.1073/pnas.0610772104, URL http://dx.doi.org/10.1073/pnas.0610772104

Ebrahim A, Lerman JA, Palsson BO, et al (2013) COBRApy: COnstraints-Based Reconstruction and Analysis for Python. BMC Syst Biol 7:74

Erdrich P, Steuer R, Klamt S (2015) An algorithm for the reduction of genome-scale metabolic network models to meaningful core models. BMC Syst Biol 9:48

Folger O, Jerby L, Frezza C, et al (2011) Predicting selective drug targets in cancer through metabolic networks. Mol Syst Biol 7:501

Foster CJ, Gopalakrishnan S, Antoniewicz MR, et al (2019) From Escherichia coli mutant 13C labeling data to a core kinetic model: A kinetic model parameterization pipeline. PLoS Comput Biol 15(9):e1007.319

Ghaffari P, Mardinoglu A, Asplund A, et al (2015a) Identifying anti-growth factors for human cancer cell lines through genome-scale metabolic modeling. Scientific Reports 5(1). 10.1038/srep08183, URL http://dx.doi.org/10.1038/srep08183

Ghaffari P, Mardinoglu A, Nielsen J (2015b) Cancer metabolism: A modeling perspective. Frontiers in Physiology 6. 10.3389/fphys.2015.00382, URL http://dx.doi.org/10.3389/fphys.2015.00382

Goldberg RN, Tewari YB, Bhat TN (2004) Thermodynamics of enzyme-catalyzed reactions–a database for quantitative biochemistry. Bioinformatics 20(16):2874–2877

Hanahan D, Weinberg RA (2011) Hallmarks of cancer: The next generation. Cell 144(5):646–674. 10.1016/j.cell.2011.02.013, URL http://dx.doi.org/10.1016/j.cell.2011.02.013

Hoops S, Sahle S, Gauges R, et al (2006) COPASI–a COmplex PAthway SImulator. Bioinformatics 22(24):3067–3074

Hosios AM, Hecht VC, Danai LV, et al (2016) Amino Acids Rather than Glucose Account for the Majority of Cell Mass in Proliferating Mammalian Cells. Dev Cell 36(5):540–549

Hucka M, Finney A, Sauro HM, et al (2003) The systems biology markup language (SBML): a medium for representation and exchange of biochemical network models. Bioinformatics 19(4):524–531

Jerby L, Shlomi T, Ruppin E (2010) Computational reconstruction of tissue-specific metabolic models: application to human liver metabolism. Molecular systems biology 6:401. 10.1038/msb.2010.56, [Online; accessed 2024-08-09]

Kacser H, Burns JA (1973) The control of flux. Symp Soc Exp Biol 27:65–104

von Kamp A, Thiele S, Hädicke O, et al (2017) Use of CellNetAnalyzer in biotechnology and metabolic engineering. J Biotechnol 261:221–228

Kanehisa M, Goto S (2000) Kegg: kyoto encyclopedia of genes and genomes. Nucleic acids research 28(1):27–30. 10.1093/nar/28.1.27, [Online; accessed 2024-08-09]

Kroll A, Rousset Y, Hu XP, et al (2023) Turnover number predictions for kinetically uncharacterized enzymes using machine and deep learning. Nature Communications 14(1):4139

König M, Bulik S, Holzhütter HG (2012) Quantifying the contribution of the liver to glucose homeostasis: A detailed kinetic model of human hepatic glucose metabolism. PLoS computational biology 8:e1002.577. https://doi.org/10.1371/journal.pcbi.1002577

Lander ES, Linton LM, Birren B, et al (2001) Initial sequencing and analysis of the human genome. Nature 409(6822):860–921. 10.1038/35057062, URL http://dx.doi.org/10.1038/35057062

Lewis JE, Forshaw TE, Boothman DA, et al (2021) Personalized genome-scale metabolic models identify targets of redox metabolism in radiation-resistant tumors. Cell Systems 12(1):68–81.e11. 10.1016/j.cels.2020.12.001, URL http://dx.doi.org/10.1016/j.cels.2020.12.001

Lewis NE, Hixson KK, Conrad TM, et al (2010) Omic data from evolved E. coli are consistent with computed optimal growth from genome-scale models. Mol Syst Biol 6:390

Li F, Yuan L, Lu H, et al (2022) Deep learning-based k cat prediction enables improved enzyme-constrained model reconstruction. Nature Catalysis 5(8):662–672

Marín-Hernández A, Gallardo-Pérez JC, Rodríguez-Enríquez S, et al (2011) Modeling cancer glycolysis. Biochimica et Biophysica Acta (BBA) Bioenergetics 1807(6):755–767. 10.1016/j.bbabio.2010.11.006, URL http://dx.doi.org/10.1016/j.bbabio.2010.11.006

Marín-Hernández A, López-Ramírez SY, Del Mazo-Monsalvo I, et al (2014) Modeling cancer glycolysis under hypoglycemia, and the role played by the differential expression of glycolytic isoforms. The FEBS Journal 281(15):3325–3345. 10.1111/febs.12864, URL http://dx.doi.org/10.1111/febs.12864

Milacic M, Beavers D, Conley P, et al (2023) The reactome pathway knowledgebase 2024. Nucleic Acids Research 52(D1):D672–D678. 10.1093/nar/gkad1025, URL http://dx.doi.org/10.1093/nar/gkad1025

Moreno-Sánchez R, Álvaro Marín-Hernández, Gallardo-Pérez JC, et al (2018) Control of the nadph supply and gsh recycling for oxidative stress management in hepatoma and liver mitochondria. Biochimica et Biophysica Acta (BBA) - Bioenergetics 1859(10):1138–1150. 10.1016/j.bbabio.2018.07.008, URL https://www.sciencedirect.com/science/article/pii/S0005272818302196

Mulukutla BC, Yongky A, Grimm S, et al (2015) Multiplicity of steady states in glycolysis and shift of metabolic state in cultured mammalian cells. PLoS One 10(3):e0121.561

Murabito E, Smallbone K, Swinton J, et al (2011) A probabilistic approach to identify putative drug targets in biochemical networks. J R Soc Interface 8(59):880–895

Murabito E, Verma M, Bekker M, et al (2014) Monte-Carlo modeling of the central carbon metabolism of Lactococcus lactis: insights into metabolic regulation. PLoS One 9(9):e106.453

Pavlova N, Thompson C (2016) The emerging hallmarks of cancer metabolism. Cell Metabolism 23(1):27–47. 10.1016/j.cmet.2015.12.006, URL http://dx.doi.org/10.1016/j.cmet.2015.12.006

Petitprez A, Poindessous V, Ouaret D, et al (2013) Acquired irinotecan resistance is accompanied by stable modifications of cell cycle dynamics independent of MSI status. Int J Oncol 42(5):1644–1653

Reder C (1988) Metabolic control theory: a structural approach. J Theor Biol 135(2):175–201

Resendis-Antonio O, González-Torres C, Jaime-Muñoz G, et al (2015) Modeling metabolism: A window toward a comprehensive interpretation of networks in cancer. Seminars in Cancer Biology 30:79–87. 10.1016/j.semcancer.2014.04.003, URL https://www.sciencedirect.com/science/article/pii/S1044579X14000510, cancer modeling and network biology

Robinson JL, Kocabaş P, Wang H, et al (2020) An atlas of human metabolism. Sci Signal 13(624)

Röhl A, Bockmayr A (2017) A mixed-integer linear programming approach to the reduction of genome-scale metabolic networks. BMC Bioinformatics 18(1):2

Roy M, Finley SD (2017) Computational Model Predicts the Effects of Targeting Cellular Metabolism in Pancreatic Cancer. Front Physiol 8:217

Schuster S, Hilgetag C (1994) On elementary flux modes in biochemical reaction systems at steady state. Biol Syst 2:165–182

Shestov AA, Liu X, Ser Z, et al (2014) Quantitative determinants of aerobic glycolysis identify flux through the enzyme GAPDH as a limiting step. Elife 3

Shlomi T, Benyamini T, Gottlieb E, et al (2011) Genome-scale metabolic modeling elucidates the role of proliferative adaptation in causing the warburg effect. PLoS computational biology 7(3):e1002.018

Singh D, Lercher MJ (2020) Network reduction methods for genome-scale metabolic models. Cell Mol Life Sci 77(3):481–488

Smith RW, van Rosmalen RP, Martins Dos Santos VAP, et al (2018) DMPy: a Python package for automated mathematical model construction of largescale metabolic systems. BMC Syst Biol 12(1):72

Snaebjornsson MT, Poeller P, Komkova D, et al (2025) Targeting aldolase a in hepatocellular carcinoma leads to imbalanced glycolysis and energy stress due to uncontrolled FBP accumulation. Nat Metab 7(2):348–366

Steuer R, Junker BH (2009) Computational Models of Metabolism: Stability and Regulation in Metabolic Networks, John Wiley & Sons, Ltd, pp 105–251. 10.1002/9780470475935.ch3, URL https://onlinelibrary.wiley.com/doi/abs/10.1002/9780470475935.ch3, https://onlinelibrary.wiley.com/doi/pdf/10.1002/9780470475935.ch3

Steuer R, Gross T, Selbig J, et al (2006) Structural kinetic modeling of metabolic networks. Proc Natl Acad Sci U S A 103(32):11,868–11,873

Teusink B, Passarge J, Reijenga CA, et al (2000) Can yeast glycolysis be understood in terms of in vitro kinetics of the constituent enzymes? testing biochemistry. European Journal of Biochemistry 267(17):5313–5329. 10.1046/j.1432-1327.2000.01527.x, URL https://febs.onlinelibrary.wiley.com/doi/abs/10.1046/j.1432-1327.2000.01527.x, https://arxiv.org/abs/ https://febs.onlinelibrary.wiley.com/doi/pdf/10.1046/j.1432-1327.2000.01527.x

Tummler K, Kühn C, Klipp E (2015) Dynamic metabolic models in context: biomass backtracking. Integr Biol (Camb) 7(8):940–951

Vander Heiden MG, DeBerardinis RJ (2017) Understanding the intersections between metabolism and cancer biology. Cell 168(4):657–669. 10.1016/j.cell.2016.12.039, URL http://dx.doi.org/10.1016/j.cell.2016.12.039

Wahl ML, Owen JA, Burd R, et al (2002) Regulation of intracellular pH in human melanoma: potential therapeutic implications. Mol Cancer Ther 1(8):617–628

Wang L, Hatzimanikatis V (2006a) Metabolic engineering under uncertainty. i: Framework development. Metabolic Engineering 8(2):133–141. 10.1016/j.ymben.2005.11.003, URL http://dx.doi.org/10.1016/j.ymben.2005.11.003

Wang L, Hatzimanikatis V (2006b) Metabolic engineering under uncertainty—ii: Analysis of yeast metabolism. Metabolic Engineering 8(2):142– 159. 10.1016/j.ymben.2005.11.002, URL http://dx.doi.org/10.1016/j.ymben.2005.11.002

Wittig U, Rey M, Weidemann A, et al (2017) SABIO-RK: an updated resource for manually curated biochemical reaction kinetics. Nucleic Acids Research 46(D1):D656–D660

Wolf J, Heinrich R (2000) Effect of cellular interaction on glycolytic oscillations in yeast: a theoretical investigation. Biochem J 345(2):321–334

Wolf J, Passarge J, Somsen OJ, et al (2000) Transduction of intracellular and intercellular dynamics in yeast glycolytic oscillations. Biophys J 78(3):1145– 1153

Yizhak K, Chaneton B, Gottlieb E, et al (2015) Modeling cancer metabolism on a genome scale. Molecular Systems Biology 11(6). 10.15252/msb.20145307, URL http://dx.doi.org/10.15252/msb.20145307

